# Sulfur isotopes in hunted ungulates reveal Palaeolithic human mobility patterns in Northern Iberia

**DOI:** 10.64898/2026.08.14.744276

**Authors:** Borja González-Rabanal, Jennifer R. Jones, Marco Vidal-Cordasco, Lucía Agudo Pérez, Adrián Álvarez-Vena, Leire Torres-Iglesias, Jesús García-Sánchez, Mónica Fernández-García, Hazel Reade, Alicia Sanz-Royo, Jeanne M. Geiling, Jesús Altuna, Koro Mariezkurrena, Maria Soledad Corchón-Rodríguez, David Cuenca-Solana, M. R. González Morales, Igor Gutiérrez-Zugasti, Rhiannon E. Stevens, Pilar Fatás, Marco de la Rasilla, Tamsin O’Connell, Michael P. Richards, Lawrence G. Straus, Ana B. Marín-Arroyo

## Abstract

This research addresses a central question in Palaeolithic research: how hunter-gatherer mobility was structured across space and time in the Cantabrian Region (northern Iberia), which has human occupation evidence spanning from the Middle Pleistocene through the Holocene. A multidisciplinary framework integrating primarily δ³⁴S isotope values, combined with δ¹³C and δ¹⁵N, palaeoproteomics, Bayesian age modelling, palaeoclimatic reconstruction, isoscape mapping, and ecological diversity was developed. A total of 905 animal bone collagen samples, with evidence of anthropogenic modifications, from 16 key archaeological sites from the Mousterian to Mesolithic (Marine Isotopic Stage 5 to 1, between 100-7 ka BP) were analysed, permitting the reconstruction of spatial patterns of resource exploitation and human mobility.

The δ³⁴S isotope values show weak, inconsistent relationships with climatic proxies, suggesting that sulfur signatures are primarily driven by geographic and ecological factors rather than climate. Strong spatial trends are observed, with higher δ³⁴S values in coastal zones and lower values inland. Diachronic trends reveal marked shifts in human mobility: smaller ranges during the Mousterian, increasing mobility through the Châtelperronian and especially the Aurignacian, followed by reduced mobility in the Gravettian and Solutrean, and renewed territorial expansion during the Magdalenian and, likely, the Azilian. In contrast, the Mesolithic is characterised by decreased mobility and thus increased territoriality in both coastal and inland contexts.

Faunal isotope values and isoscape predictions reveal that some animals were acquired beyond local foraging ranges during the Palaeolithic, particularly in inland regions with lower δ³⁴S values. Isotopic niche analyses indicate partial interspecific overlap consistent with ecological flexibility. Macromammal and micromammal diversity exhibit contrasting patterns, with a significant negative correlation in Simpson and Shannon indices. Macromammal diversity correlates negatively with δ³⁴S values, linking increased hunting diversity to expanded catchment areas and longer-distance foraging, whereas micromammal diversity shows positive correlations with δ³⁴S, δ¹³C and δ¹⁵N reflecting stronger climatic influence.

Overall, these results demonstrate that hunter-gatherer behaviour in northern Iberia during the Middle and Late Palaeolithic was highly dynamic, combining logistical and residential strategies that shifted in response to changing environmental conditions, resource distributions and cultural adaptations.

## Introduction

Hunting is recognised as one of the earliest economic behaviour in human evolution and was the primary source of subsistence in Western Eurasia until the advent of agriculture and livestock domestication 8,000 years ago (Straus, 1987; Finlayson, 2017; Carbonell and Canals-Salomó, 2026). The nomadic lifestyle of those hunter-gatherers meant moving regularly to survive. The frequency and distance of foragers’ movements were multifaceted, depending on factors such as latitude, geography, natural barriers, soil, climate, resource availability, and human population size (Kelly, 1981; Binford, 1982). Hunter-gatherers used to make short logistical and/or residential moves within very constrained territorial boundaries, especially in areas of abundant resources and an equable climate, but also undertake enormous annual rounds in harsh environments (Speth, 2022).

The Cantabrian Region is located on the Atlantic coast of northern Iberia (Fig. 1A). It consists of a narrow corridor, ∼400 km long and 35 to 60 km wide, with an Atlantic climate characterised by temperate, wet conditions, belonging to the Eurosiberian biogeographic region (Rivas-Martínez et al., 2004). It is a region geographically separated from the Spanish central plateau and the Ebro valley by steep, high mountains (the Cantabrian Cordillera) that run parallel to the coast and reach heights of over 2500 m above present sea level. This high-relief coastal strip is characterised by generally short, steep river valleys that descend mostly perpendicularly, from south to north, into the Cantabrian Sea (also known as the Bay of Biscay) (García Codron, 2004). Thus, the special orography of this area conditioned the adaptations and patterns of exploitation of northern Iberian landscapes by Stone Age hunter-gatherers, allowing them to utilise a wide variety of closely spaced habitats in which to develop their economic strategies from the shore to high mountain slopes, along the axes of numerous river valleys (Straus, 1992; González Sainz, 2003). The Cantabrian Region is a key area for disentangling the economic transformations of hunter-gatherer adaptive systems, as it preserves many relevant Palaeolithic sites with high-resolution chronologies, along with an abundant faunal record related to human diet that serves as a biomolecular framework for unravelling diachronic human hunting behaviour (Marín-Arroyo et al., 2018).

**Fig. 1.**
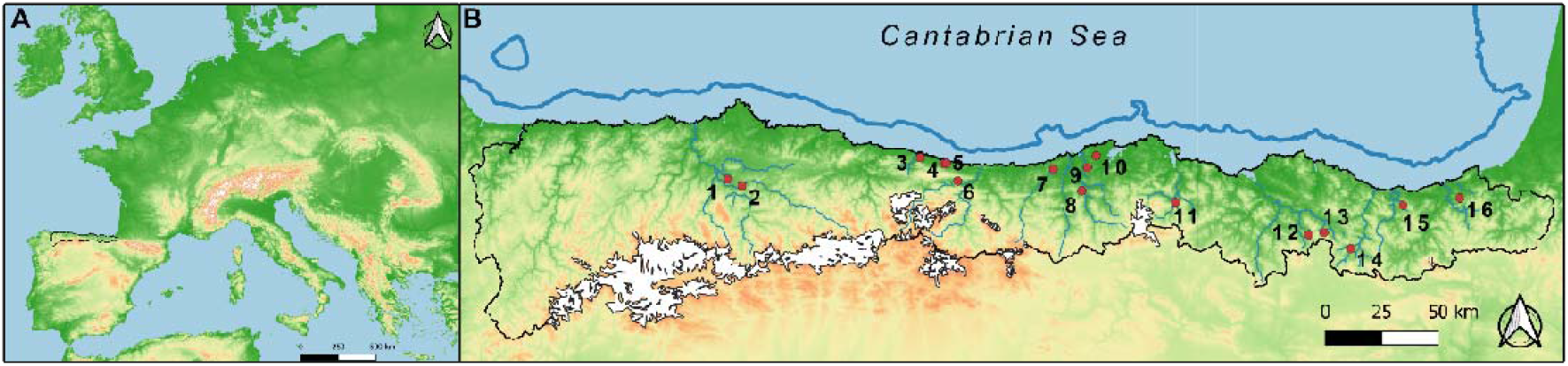
A) Geographical location of the Cantabrian Region in Western Europe. B) Location of the archaeological sites studied in this work: 1. Las Caldas; 2. La Viña; 3. La Riera; 4. El Toral III; 5. El Mazo; 6. Llonín; 7. Altamira; 8. El Castillo; 9. Covalejos; 10. El Juyo; 11. El Mirón; 12. Axlor; 13. Bolinkoba; 14. Labeko Koba; 15. Amalda; 16. Aitzbitarte III. White polygons show the maximum extension of the Cantabrian Mountain glaciers (Rodríguez-Rodríguez et al., 2015) and the blue line indicates the furthest coastline during the Late Glacial Maximum (Bilbao-Lasa et al., 2020).

Almost half a century after Lewis Binford’s publication (1980) of “*Willow Smoke and Dogs’ Tails: Hunter-Gatherer Settlement Systems and Archaeological Site Formation”,* the forager-collector model remains a fundamental milestone in the study of hunter-gatherer settlement and land use, establishing the distinction between residential and logistical mobility. In the 1980s, following his approach, K.W. Butzer (1986) and L.G. Straus (1986) proposed complementary models for Palaeolithic adaptations and settlement in the Cantabrian Region, highlighting the distinction between coastal and montane occupations. Butzer’s Model A proposed that hunter-gatherers situate their base camps along the piedmont belt, from which individual bands could exploit the coastal area, lower and upper valleys, and Cordilleran slopes. Model B, in contrast, formulated separate coastal zones and montane band territories, practising different subsistence strategies (Butzer, 1986). On the other hand, Straus’ model postulated the existence of residential base camps, mainly located on the coastal plain, from which logistical trips would be made to exploit resources along the shore and in the montane range (Straus, 1986), as later demonstrated by Marín-Arroyo (2009a; 2009b) by using optimal foraging theory and resource catchment areas estimation. Nonetheless, the cultural territories of these hunter-gatherer groups would have extended beyond the area where immediate subsistence resources were available, which would help explain the technological and artistic similarities found along the northern Atlantic coast, often notably extending to SW France, the French Pyrenees and beyond. This is evidenced in the circulation of raw materials, marine mammal remains, molluscs, certain highly distinctive osseous artefacts and symbolic motifs in both rock and portable art objects (Straus, 2005; Sauvet et al., 2008; Rivero, 2010; Straus, 2018a; Lefebvre et al., 2021).

Quaternary climate in the Cantabrian Region was characterised by alternating cold and warm cycles (Sanchez Goñi and Harrison, 2010). MIS 3 (59.4-27.8 ka BP) was a climatically unstable period marked by rapid millennial-scale oscillations between cold stadials and temperate interstadials (d’Errico and Sánchez Goñi, 2003). MIS 2 (27.8–14.7 ka BP) corresponds to a cold, arid phase with expanded open landscapes and glacier advances (Moreno et al., 2012), including the Last Glacial Maximum (LGM), when continental ice sheets reached their maximum extent and sea levels dropped by ∼130 m (Clark et al., 2009; Lambeck et al., 2014). However, the Cantabrian montane glaciers reached their maximum extent during MIS 3 between 48-35 ka BP (Serrano et al., 2017). MIS 1 (14.7 ka BP to present) marks a major warming phase beginning with the Bølling–Allerød interstadial (14.6-12.9 ka BP) followed by the Younger Dryas cold event (12.9–11.7 ka BP) (Sánchez-Morales, 2023). The Pleistocene/Holocene transition brought significant environmental transformations that led to changes in the subsistence strategies and settlement patterns of prehistoric human groups of the Cantabrian Region (Altuna, 1990; Marín-Arroyo, 2013). Climate shifts led to increased temperatures (albeit with temporary downturns), ultimately causing the final melting of montane glaciers, sea-level rise, and coastal transgression (Straus, 2018b). At the same time, humidity and rainfall increased, leading to the expansion of mixed deciduous forests from the glacial-age open landscapes with few trees (often mainly scattered pines and birches) (Sánchez-Morales, 2023). Flora and ecologically flexible faunal species such as red deer became adapted to more temperate climatic conditions, whilst narrowly cold-adapted species such as mammoth, woolly rhinoceros or reindeer disappeared (Altuna, 1992; Álvarez-Lao and García, 2010).

The archaeozoological research indicates that red deer was the most abundant and exploited prey in the Cantabrian Region during the Middle/Upper Palaeolithic transition (Altuna, 1988; 1989; Altuna and Mariezkurrena, 1988; 2020; Yravedra-Sainz de los Terreros et al., 2015; Marín-Arroyo and Sanz-Royo, 2022), the Late Upper Palaeolithic technocomplexes (Altuna, 1972; 1990; 1994; Freeman, 1973; Straus, 1977; Portero et al., 204; 2026; Geiling et al., 2025), and the last Azilian and Mesolithic hunter-gatherers cultures (Altuna, 1979; 1998; Marín-Arroyo, 2013; Portero et al., 2022). This game animal was locally supplemented (or even outnumbered) by ibex (*Capra pyrenaica*) and, less frequently, chamois (*Rupicapra rupicapra*), together with small numbers of large bovids (*Bos primigenius* and *Bison priscus*) and horses (*Equus ferus*), with fluctuating percentages of species representation depending on the site location and the chronocultural period analysed (Altuna, 1972; Marín-Arroyo, 2009b). The available Azilian and Mesolithic archaeozoological records reveal a shift in hunting preferences, which are now more diversified, including ungulates scarcely present or exploited before, such as wild boar (*Sus scrofa)* and roe deer (*Capreolus capreolus*), to the still-dominant red deer (Altuna, 1998; Fano, 2019), but with an increase in the consumption of juvenile prey (Marín-Arroyo, 2013) and the intensification in the exploitation of marine resources (Álvarez-Fernández, 2011; Gutiérrez-Zugasti, 2011; Arniz-Mateos et al., 2024; García-Escárzaga et al., 2024).

Sulfur-stable isotope measurements in bone collagen have demonstrated their potential to track the geographical origins of humans and animals recovered from archaeological sites, given the influence of local geology, water cycles and proximity to the sea on the sulfur cycle (Richards, 2023; Stevens et al., 2025). Sulfur shows a low trophic enrichment (± 0.5 ± 2.4 ‰) (Krajcarz et al., 2019) within the biosphere, representing a long-term dietary average over the last decade of a living individual (Richards et al., 2001). However, it is less abundant in the pool of amino acids than carbon and nitrogen, so it may reflect a much longer average lifespan of the individual (Thode, 1991). As sulfur isotopic values are directly linked to local geology, soil type, proximity to the sea and rainfall (Nehlich, 2015), terrestrial ecosystems have a very variable δ³⁴S values (from -20‰ to +30‰), reflecting isotope values of the local geological sulfate, which are influenced by weathering of soils from different rock types which affects sulfur bioavailability (Krouse et al., 1996; Brenot et al., 2007). Marine environments exhibit a more homogeneous signature, with values close to 20‰. Oceanic sulfur is redeposited as rain over coastal platforms, reaching distances of up to 50 km offshore due to the sea-spray effect (Wadleigh et al., 1996). Greater variability in sulfur isotope values is observed in freshwater landscapes, ranging from - 22‰ to 20‰. This wide range is in part due to sulfate-reducing microbes living in rivers or lakes. Recently, lower and very negative sulfur values have been linked to changing permafrost conditions (Stevens et al., 2023) and wetlands (Lamb et al., 2023; Wexler and Stevens, 2025), which may alter soil δ^34^S values due to sulfur cycling being influenced by weathering and hydrology. The low fractionation of sulfur within the food chain means that animal δ^34^S values reflect bioavailable δ^34^S from primary producers at the bottom of the food chain, and can reflect the locations where animals are feeding (Richards et al., 2003).

Faunal δ³⁴S values measured in bone collagen represent an average dietary signal of the animals’ habitual foraging areas over an extended period, which may encompass multiple landscapes depending on the species ecology, seasonal and/or migratory movements, and ethological behaviours (Nehlich, 2015). This temporal and spatial integration means that faunal sulfur isotope values should be interpreted with caution, as they reflect broad patterns of landscape use rather than a single location (Stevens et al., 2025). The occurrence within the same archaeological site or stratigraphic level of faunal remains showing clear evidence of anthropogenic modifications in association with other evidence of human activities (i.e. hearths, stone tools, etc.), which display distinct δ³⁴S values, suggests that these ungulates likely exploited different isotopic landscapes throughout the period represented by bone collagen (Jones et al., 2018). Consequently, as hunting decisions are shaped by prey ecology, environmental conditions, and human subsistence strategies, the sulfur isotopic variability among anthropogenically transported and accumulated faunal remains can provide valuable information on the diversity of hunting range exploited by hunter-gatherers and, ultimately, contribute to reconstructing patterns of hunter-gatherer mobility and landscape use (Richards, 2023).

In recent years, sulfur research in Late Pleistocene mammal assemblages has revealed its potential to infer differences in human hunting range across space and time (Jones et al., 2018; 2019; Wißing et al., 2019; Britton et al., 2023b; 2023a; Pederzani et al., 2023; Barakat et al., 2026), but it also has allowed the identification of environmental and climatic changes at a local scale (Drucker et al., 2011; Reade et al., 2020; 2021; Stevens et al., 2023). However, given the equifinality in sulfur isotopes, understanding the exploitation of the landscape and the mobility behaviour of past human groups remains complex and poorly understood (Richards, 2023). Similarly, spatial modelling of sulfur data is still in its early stages (Bataille et al., 2021; González-Rabanal et al., 2025).

Here, we present the largest and most temporally extensive sulfur isotope dataset (n=905) for a European Palaeolithic context, encompassing archaeological levels from the Mousterian to the Mesolithic and spanning Marine Isotope Stages (MIS) 5 to 1, from 100-7 ka BP. The aim of this work is to investigate patterns of δ^34^S trends for ungulate taxa, climatic phases, and cultural contexts documented in the Cantabrian Region. By integrating this dataset, we aim to reconstruct a high-resolution Palaeolithic sulfur isoscape, providing a robust framework for evaluating mobility patterns and hunting territories of Palaeolithic hunter-gatherers over time in northern Iberia. To accomplish this goal, we adopt an innovative and multidisciplinary approach that combines stable isotopes with paleoproteomics, Bayesian chronological modelling, palaeoclimatic reconstructions, isoscape modelling, catchment area estimation and ecological diversity analysis.

## Methods

### Materials

Bone collagen samples from 16 Cantabrian sites with well-established stratigraphical and chronological sequences were analysed for δ^13^C, δ^15^N and δ^34^S using IRMS (Table 1). From west to east, these sites are located in the modern-day provinces of Asturias (Las Caldas, La Viña, La Riera, El Toral III, El Mazo and Llonín), Cantabria (Altamira, El Castillo, Covalejos, El Juyo and El Mirón), Vizcaya (Axlor and Bolinkoba) and Guipúzcoa (Labeko Koba, Amalda and Aitzbitarte III) (Fig. 1B; SText 1). The chronometric measurements of these sites cover a time range from 100,000 to 7,000 ka BP, deriving from Mousterian (*n*= 6), Châtelperronian (*n*= 1), Aurignacian (*n*= 5), Gravettian (*n*= 6), Solutrean (*n*= 5), Magdalenian (*n*= 6), Azilian (*n*= 2) and Mesolithic (*n*= 4) cultural technocomplexes (STable 1). All faunal specimens are mammals and come from 111 archaeological levels (Table 2, SText 1; STable 2). From a paleoclimatic point of view, these archaeological levels are framed during the MIS 5 to MIS 1: MIS 5 (130-73.5 ka BP) (*n*= 2), MIS 4 (73.5-59.4 ka BP) (*n*= 2), MIS 3 (59.4-27.8 ka BP) (*n*= 36), MIS 2 (27.8–14.7 ka BP) (*n*= 55), and MIS 1 (14.7 ka BP-today) (*n*= 16). Detailed information about the sites is included in the S1 Text.

**Table 1.** Archaeological sites studied in this research and origin of the samples from archaeological excavations and archaeozoological studies.

| Sites | Location | Archaeological excavation | Archaeological and archaeozoological references |  |
| --- | --- | --- | --- | --- |
| Aitzbitarte III | Errentería (Guipúzcoa) | J. Altuna | Altuna et al. 2011; 2017 | Altuna and Mariezkurrena 2011; 2017 |
| Altamira | Santilla del Mar (Cantabria) | L.G. Freeman and J. González-Echegaray | Freeman and González Echegaray 2001 | Castaños Ugarte and Castaños de la Fuente 2014 |
| Amalda | Zestoa (Guipúzcoa) | J. Altuna | Altuna et al. 1990 | Altuna 1990 |
| Axlor | Dima (Vizcaya) | I. Barandiarán | Barandiarán 1980 | Altuna 1989 |
| Bolinkoba | Abadiño (Vizcaya) | I. Barandiarán | Barandiarán 1950 | Castaños and Castaños 2015 |
| Covalejos | Piélagos (Cantabria) | R. Montes and J. Sanguino | Sanguino and Montes 2005 | Yravedra-Sainz de los Terreros et al. 2015 |
| El Castillo | Puente Viesgo (Cantabria) | V. Cabrera Valdés and F. Bernaldo de Quirós | Cabrera Valdés 1984 | Luret 2017; Castaños Ugarte 2018 |
| El Juyo | Camargo (Cantabria) | L.G. Freeman and J. González-Echegaray | Freeman and González Echegaray 1981 | Klein and Cruz-Urbe 1985 |
| El Mazo | Llanes (Asturias) | I. Gutiérrez-Zugasti and D. Cuenca-Solana | Gutiérrez-Zugasti et al. 2018 | Marín-Arroyo et al. 2020 |
| El Mirón | Ramales de la Victoria (Cantabria) | L.G. Straus and M.R. González Morales | Straus and González Morales 2012 | Marín-Arroyo et al. 2023 |
| El Toral III | Llanes (Asturias) | M. Noval and I. Gutiérrez-Zugasti | Noval 2014 | Andreu Alarcón 2013 |
| La Riera | Llanes (Asturias) | L.G. Straus and G.A. Clark | Straus and Clark 1986 | Altuna 1986 |
| La Viña | Oviedo (Asturias) | J. Fortea | Fortea 1992; Rasilla et al 2018 | Rasilla et al. 2020; Torres-Iglesias 2023; Sanz-Royo 2023 |
| Labeko Koba | Arrasate (Guipúzcoa) | A. Arrizabalaga and J. Altuna | Arrizabalaga and Altuna 2000 | Altuna and Mariezkurrena 2000 |
| Las Caldas | San Juan de Priorio (Asturias) | M.S. Corchón Rodríguez | Corchón Rodríguez 2017a; 2017b | Altuna and Mariezkurrena 2017a; 2017b |
| Llonín | Peñamellera Alta (Asturias) | J. Fortea | Fortea et al. 1992; 1995; 1999; Rasilla et al. 2014 | Sanchís et al. 2019; Rasilla et al. 2020 |

**Table 2.**
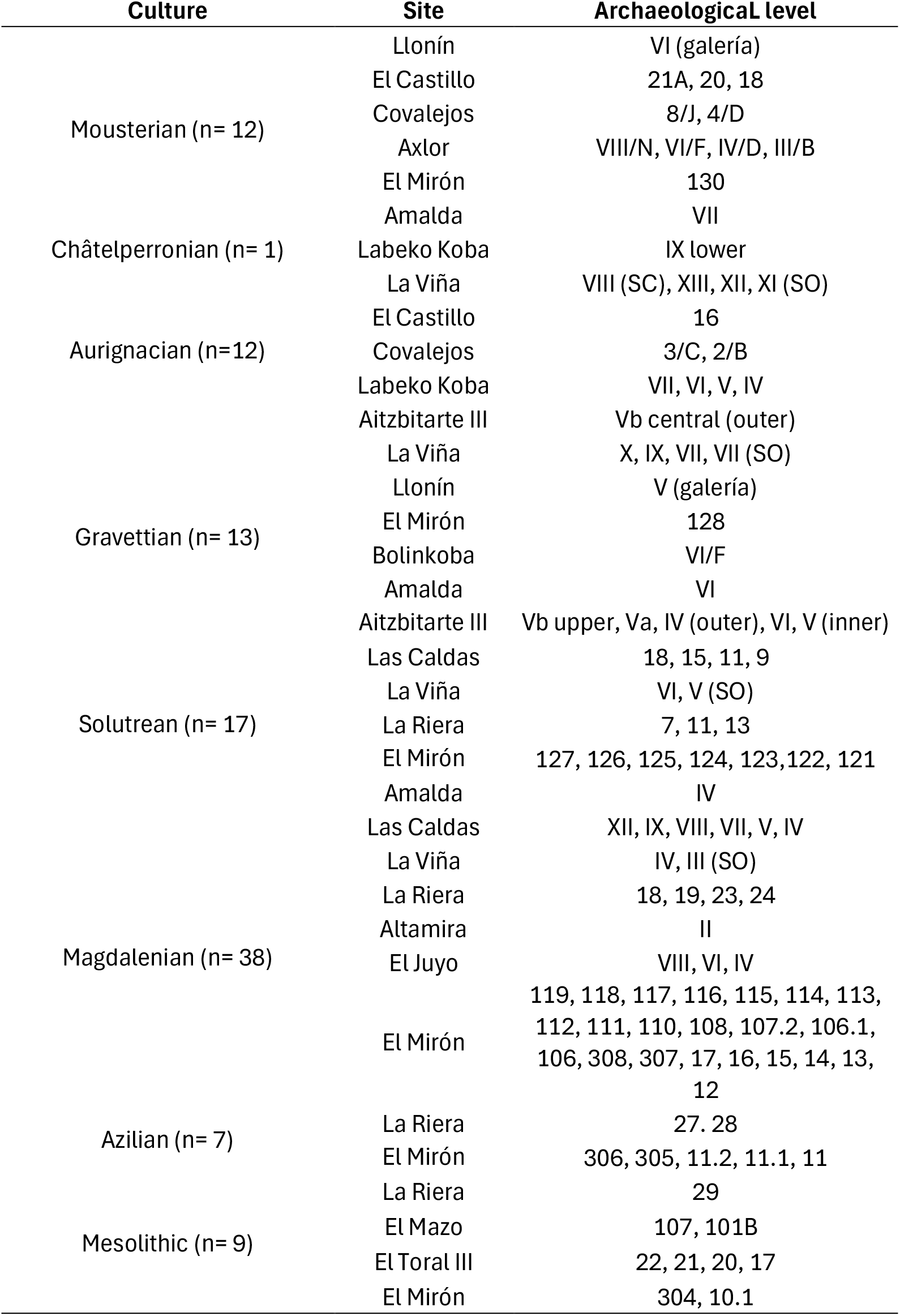
Archaeological layers analysed in this study by culture and site. Levels 19 and 18 from El Castillo are considered archaeologically sterile.

The most commonly hunted mammal species during the Late Pleistocene at a regional scale were sampled, including four main taxa: red deer (*Cervus elaphus*) (*n*= 483), ibex (*Capra pyrenaica)* (*n=* 196), horse (*Equus ferus*) (*n*= 107), aurochs/bison (*Bos/Bison* sp.) (*n*= 100); plus other additional species, such as wild boar (*Sus scrofa*) (*n*= 7), roe deer (*Capreolus capreolus*) (*n*= 6), and reindeer (*Rangifer tarandus*) (*n*= 6). The selection of the faunal samples in museum collections was based on the following criteria: 1) stratigraphic position of the remains at the site; 2) selection of bones labelled to assure their provenance within the site; 3) selection of adult animal bones taxonomically and anatomically identifiable and 4) selection of anatomical elements with evident anthropogenic modifications (i.e., cut marks and/or percussion marks) associated to lithic industries. Sample information for each specimen, including site, archaeological level, culture, sample number, animal species, and anatomical element, is provided in STable 3.

In total, 905 faunal specimens have been analysed in this work (Table 3), including 799 unpublished and 106 published δ^34^S values. Additionally, 438 unpublished and 467 published δ^13^C and δ^15^N values were used to explore comparisons and correlations with δ^34^S values (Stevens et al., 2014; Jones et al., 2018; 2019; 2020; 2021; Pederzani et al., 2023). The δ^34^S values were generated from the same collagen samples extracted for these studies, or, if insufficient collagen was available, the same bone fragments were resampled to obtain new collagen samples.

**Table 3.** Bone collagen samples by site and species analysed for δ^13^C, δ^15^N and δ^34^S in this research.

| Sites | *Bos/Bison* sp. | *Capra pyrenaica* | *Capreolus capreolus* | *Cervus elaphus* | *Equus ferus* | *Rangifer tarandus* | *Sus scrofa* | Total |
| --- | --- | --- | --- | --- | --- | --- | --- | --- |
| Aitzbitarte III | 14 |  |  | 16 | 5 |  |  | 35 |
| Altamira |  |  |  | 9 |  |  |  | 9 |
| Amalda | 5 | 7 |  | 11 | 14 |  |  | 37 |
| Axlor | 26 |  |  | 35 | 4 |  |  | 65 |
| Bolinkoba | 5 |  |  | 4 | 7 | 1 |  | 17 |
| Covalejos | 6 |  |  | 31 | 2 |  |  | 39 |
| El Castillo | 22 |  |  | 83 |  |  |  | 105 |
| El Juyo | 1 |  |  | 15 |  |  |  | 16 |
| El Mazo |  |  | 2 | 3 |  |  | 3 | 8 |
| El Mirón |  | 99 |  | 79 |  |  |  | 178 |
| El Toral III |  |  | 4 | 15 |  |  | 4 | 23 |
| La Riera |  | 42 |  | 56 |  | 2 |  | 100 |
| La Viña |  | 15 |  | 54 | 23 | 3 |  | 95 |
| Labeko Koba | 21 |  |  | 15 | 10 |  |  | 46 |
| Las Caldas |  | 33 |  | 50 | 42 |  |  | 125 |
| Llonín |  |  |  | 7 |  |  |  | 7 |
| **Total** | **100** | **196** | **6** | **483** | **107** | **6** | **7** | **905** |

### Stable isotope analysis of bone collagen

Faunal specimens were sampled by the EvoAdapta Research Group from the collections deposited at the Museo Arqueológico de Asturias (MAA), the Museo de Prehistoria y Arqueología de Cantabria (MUPAC), the Museo de Arqueología de Bizkaia (Arkeologi Museoa) and the Centro de Colecciones Patrimoniales de la Diputación Foral de Gipuzkoa (Gordailua Center), where the materials are currently curated. Sampling was based on extracting ∼1 g of bone using a low-vibration micromotor with a diamond-edge cutting wheel in dedicated clean spaces, in accordance with the protocols of these institutions. For the majority of the newly generated δ^34^S values within this paper, sample preparation was performed at the University of Cantabria using the EvoAdapta Group facilities, where bone collagen extraction was undertaken according to the procedures proposed by Richards and Hedges (1999). This method involves the following steps: 1) cleaning of the bone fragments (0.6*–*0.8 g) by mechanical abrasion to remove any possible contamination; 2) demineralisation of the samples in 0.5 M HCl at 6*–*8 °C, typically between 3*–*10 days; 3) washing using de-ionized water; 4) gelatinisation of the samples in a weak solution of pH 3 HCL at 70 °C for 48 h; 5) filtration with 5*–*8 μm Ezee® filters; 6) freeze-drying of the samples. Samples were analysed for δ^13^C, δ^15^N and δ^34^S using a Europa ScientificTM elemental analyser coupled to a mass spectrometer (EA-IRMS) at the former Iso-Analytical laboratory (now known as Isolab) in the United Kingdom. Samples with S-EVA, S-UBC and XLHB codes were prepared and analysed for δ^13^C and δ^15^N at the Max Planck Institute for Evolutionary Anthropology, the University of British Columbia, and the University of Cambridge, respectively, following the same methodology, with an added ultrafiltration step.

The δ^13^C, δ^15^N, and δ^34^S values were reported relative to the V-PDB, AIR, and VCDT international standards, and are shown in STable 3. The reference material used for carbon and nitrogen isotope analysis of the collagen samples was IA-R068 (soy protein, δ^13^C= -25.22‰, δ^15^N= 0.99‰). IA-R068, IA-R038 (L-alanine, δ13C= -24.99‰, δ^15^N= -0.65‰), IA-R069 (tuna protein, δ^13^C= -18.88‰, δ^15^N= 11.60‰) and a mixture of IAEA-C7 (oxalic acid, δ^13^C= -14.48‰) and IA-R046 (ammonium sulfate, δ^15^N= 22.04‰) were run as quality control check standards. IA-R068, IA-R038 and IA-R069 are calibrated against and traceable to IAEA-CH-6 (sucrose, δ^13^C= -10.45‰) and IAEA-N-1 (ammonium sulfate, δ^15^N= 0.40‰). IA-R046 is calibrated against and traceable to IAEA-N-1. IAEA-C7, IAEA-CH-6 and IAEA-N-1 are interlaboratory comparison standards distributed by the International Atomic Energy Agency, Vienna. The reference material used for sulfur isotope analysis of the collagen samples was IA-R061 (barium sulfate, δ^34^S= 20.33‰). IA-R061, IA-R025 (barium sulfate, δ^34^S= 8.53‰) and IA-R026 (silver sulfide, δ^34^S= 3.96‰) were used for calibration and correction of the ^18^O contribution to the SO+ ion beam. IA-R061, IA-R025 and IA-R026 are in-house standards calibrated against and traceable to NBS-127 (barium sulfate, δ^34^S= 20.3‰) and IAEA-S-1 (silver sulfide, δ^34^S= -0.30‰). IA-R061, IAEA-S-1, IA-R068 (soy protein, δ^34^S= 5.25‰) and IA-R069 (tuna protein, δ^34^S= 18.91‰) were measured as quality control check standards during the batch analysis of the collagen samples. IA-R068 and IA-R069 are in-house standards calibrated against and traceable to NBS-127 and IAEA-SO-5 (barium sulfate, δ^34^S= 0.50‰). NBS-127, IAEA-S-1 and IAEA-SO-5 are inter-laboratory comparison standards distributed by the International Atomic Energy Agency (IAEA) with internationally accepted δ^34^S values. One in every five samples was measured in duplicate, and replication was typically <0.1‰, indicating high precision in the acquired data. Quality indicators habitually established were used: %Col (>1), %C (30*–*44%), %N (11*–*16%), %S (0.15*–*0.35%), C:N (2,9*–*3,6), C:S (600 ± 300) and N:S (200 ± 100) (DeNiro, 1985; Ambrose, 1990; van Klinken, 1999; Nehlich and Richards, 2009).

All statistical tests were conducted in R software (R Core Team, 2020) and the ggplot2 package was used for visualisation (Wickham, 2016). For each isotope, we compare the chronological evolution of each species values and the existing differences between species within different levels/cultures. At a site level, a Shapiro–Wilk/Kolmogorov-Smirnov test was used to confirm that all stable isotope data (δ^13^C, δ^15^N, and δ^34^S) were not normally distributed (p-value <0.05). Depending on the results, Pearson/Spearman correlation tests were used to analyse significant relationships among δ^34^S and δ^13^C/δ^15^N isotopes. Stable isotope value groupings were analysed statistically using a Wilcoxon-Mann-Whitney U test with a post-hoc Holm-Bonferroni correction. A p-value of <0.05 was deemed to be statistically different.

To evaluate the relationship between δ^34^S and other independent variables (e.g., climate, ecological diversity), we used the mean sulfur isotope value for each archaeological level as an observation in linear mixed models (LMMs). Because the same site may have multiple observations (i.e. δ^34^S values) across different archaeological units, observations are not independent, and spatial autocorrelation was assessed using Moran’s I test. We computed the *k* nearest neighbours for each observation based on spatial coordinates, converting these lists into a spatial weight matrix which was then used in a Moran’s I test. Moran’s I value ranges between 1 and - 1, with positive values suggesting spatial autocorrelation characterised by clustering, while negative values indicate negative spatial autocorrelation characterised by segregation. When p-values were significant, we fitted LMMs that incorporated spatial structure to account for spatial autocorrelation and non-independence among observations. When Moran’s test was not significant, we ran LMMs without spatial structure.

### Palaeoproteomic analysis with ZooMS

Zooarchaeology by Mass Spectrometry (ZooMS) was performed on 31 samples that had been previously taxonomically identified through archaeozoological analyses (STable 4). These were selected because their stable isotope values fell outside their taxonomic groupings. ZooMS was applied to test whether these specimens were simple outliers or, conversely, belonged to other species, with the aim of improving the reconstruction of carbon, nitrogen, and sulfur isotopic values for environmental and animal behaviour.

ZooMS analysis followed published protocols (Buckley et al., 2009; Welker et al., 2015; Reade et al., 2021) with sample preparation undertaken in the EvoAdapta laboratory. 19 samples were analysed by extracting and digesting collagen from a small amount of bone (∼10–30 mg). Meanwhile, for the remaining 12 samples, ZooMS pretreatment was performed on the collagen extracted for stable isotope analysis. Bone samples were first demineralised in 150 µL of 0.6 M hydrochloric acid at 4 °C for 18 h. After neutralising the samples with 50 mM ammonium-bicarbonate (NH_4_HCO_3_, AmBic), bone collagen was extracted by incubation at 65° C for 1 h in 100 μL of AmBic buffer. Then, 50 µl of the resulting supernatant was digested with trypsin (0.5 μg/μL, Promega) between 12 and 18 h at 37 °C. Digestion was stopped by adding 1 μL of 10% trifluoracetic acid (TFA), and the collagen peptides were concentrated and cleaned using C18 Ziptips (ThermoFisher Scientific). For samples analysed with already extracted collagen, first, a small amount of the dry collagen was collected by introducing a tip into the tube. Then, collagen was dissolved in 100 µL of AmBic, and a 50 µL aliquot of this solution was transferred to a new tube. Digestion and peptide extraction followed the same steps described above for the demineralised bone samples.

Collagen peptides were then spotted in triplicate on a Bruker MALDI plate using α-cyano-4-hydroxycinnamic acid as the matrix. MALDI-TOF MS analysis was conducted at the University of York (UK) on a Bruker UltrafleXtreme instrument, with a mass range of 800–4000 Da. Spectral triplicates for each sample were merged in R (R Core Team, 2020) using the MALDIquant package (Gibb and Strimmer, 2012) and settings previously described (Ruebens et al., 2023). Taxonomic identification was made manually in mMass v.5.5.0 (Strohalm et al., 2008), comparing the spectra against a reference database of known collagen peptide marker masses for the mammal species occurring in Europe during the Pleistocene (Buckley et al., 2009; Welker et al., 2016; Jensen et al., 2020).

### Chronological assignment

After reviewing the chronostratigraphic sequence of the 16 archaeological sites individually, 199 published chronological dates were compiled from the literature from the 111 archaeological levels with stable isotope analysis in this study (STable 1). Preferably, radiocarbon dates obtained by ultrafiltration pre-treatment method (AMS-UF) were chosen. Accelerator Mass Spectrometry (AMS) and/or conventional 14C dates have been included at levels where this protocol was not applied. Additionally, for the oldest levels, where the measurements were beyond the radiocarbon limit (∼55 ka BP), Optically Stimulated Luminescence (OSL) dates were included. To assign a chronological age to a particular archaeological level, we conducted Bayesian modelling at each site using OxCal 4.4 (Bronk Ramsey, 1995) and the INTCAL20 calibration curve (Reimer et al., 2020). For those purposes, we considered previous Bayesian age models (Wood et al., 2014; 2018; Marín-Arroyo et al., 2018; 2026; Hopkins et al., 2021; Jones et al., 2021; García-Escárzaga et al., 2022; Fernández-García et al., 2023; Arniz-Mateos et al., 2024) and new models performed *ex profeso* for this study using the same software (Bronk Ramsey, 2009). For those levels (*n*= 18 – 16% of the total) that do not have available dates, the dates of the adjacent infra or supra levels have been compiled and used to model the most probable dates of these levels (SText 2; STable 2).

### Isotope niche partitioning

Feeding and spatial niche partitioning between species were evaluated using the SIBER R package (Jackson et al., 2011) by plotting the δ^13^C-δ^15^N and δ^15^N-δ^34^S isotopic spaces as bivariate plots. The degree of overlap between the isotopic areas of each species indicates whether they occupy similar or different feeding niches. We assessed the isotopic niches of *Bos/Bison* sp., *Capra pyrenaica*, *Cervus elaphus*, *Equus ferus*, and *Rangifer tarandus* at each archaeological level, when more than three samples for a given species were available.

To estimate overlaps between niches, we used two metrics: the convex hull total area (TA), which encompasses all samples for a given species, and the standard ellipse area corrected for small sample size (SEAc), corresponding to the core area (40% of the data) (Jackson et al., 2011). We classified the degree of overlap between isotopic niche into: null overlap (0%), low overlap (< 30%), moderate overlap (30–60%), and high overlap (> 60%) (Schwartz-Narbonne et al., 2019).

### Climatic correlations

We have used two different climatic proxies to correlate the δ^34^S values from each archaeological level with climate fluctuations (STable 5). We calculated the local Mean Annual Temperature (MAT) in °C and Mean Annual Precipitation (MAP) in mm/yr for each site and level using the HadCM3B-M2.1 coupled general circulation model with active atmosphere, ocean, and sea ice components, forced by orbital parameters, greenhouse gases, and ice sheets (Krapp et al., 2021). With the aim of reconstructing the local climatic conditions under which the human groups that inhabited each occupational level of these sites, we also estimated the MAT and MAP parameters using the Bioclimatic Model (BM), a qualitative method based on the association of small-mammal communities to different climatic zones developed by Hernández Fernández (2001; 2007) and revisited by Royer et al. (2020).

Following the review of the BM method initiated by Domínguez-García et al. (2023), we have modified the Climatic Restriction Indices (CRI) for *Arvicola sapidus*, as well as for *Microtus pyrenaicus* (=*gerbii*) and *Microtus lusitanicus*, by considering the current geographic distribution provided by Wilson et al. (2017). Due to the challenges involved in accurately identifying the main genera of the Eulipotyphla present in the region (*Sorex*, *Crocidura*, *Neomys*, and *Talpa*; see Álvarez-Vena et al. (2021; 2023) for further details), we have chosen to use the BM approach based on the order Rodentia. For taxa identified only at the genus or group level, such as *Arvicola* sp. or *Microtus* (*Terricola*) ex gr. *duodecimcostatus,* respectively, we have resorted to using “chimeric” or average species, following Royer et al. (2020). For that purpose, we compiled 91 micromammal records from the literature by site and level following a pilot study on the Middle/Upper Palaeolithic transition of the Cantabrian Region (Fernández-García et al., 2023), and expanded the dataset to include all the chronological ranges involved in this work until the Mesolithic (STable 6). Small-mammal associations with a single rodent species have been excluded. Levels with a Minimum Number of Individuals (MNI) <10 were also excluded. Finally, the current MAT and MAP for each site location were estimated using the WorldClim dataset (Fick and Hijmans, 2017) (STable 7) and used to perform a delta bias correction on past MAT and MAP estimates derived from the general circulation model (Krapp et al., 2021).

### Sulfur isoscape and catchment areas

To compare sulfur isotope values measured in animal bone collagen with the bioavailable sulfur isotope values in the areas surrounding the sites under study, we combine sulfur isoscape modelling with catchment-area calculations. To predict the sulfur isotope values around the sites under study, we used the modern Iberian δ^34^S isoscape published by González-Rabanal et al. (2025), which was adapted from the European sulfur isoscape published by Bataille et al. (2021). This new isoscape used 393 sulfur isotope values from 41 sites across Iberia and placed special focus on the bioavailability of δ^34^S in the Holocene landscape of the Cantabrian Region. After incorporating 22 independent variables (geology, elevation, climate, soil properties, aerosol deposition, and proximity to the coast) in the region of interest, and applying the Random Forest (RF) machine learning algorithm, a predictive model of δ^34^S was generated. In that model, four variables, including elevation above the sea level, Bouguer anomaly, distance from the coast, and strontium content of the soil, were determined to be the dominant predictors of the δ^34^S isotope values. The final δ^34^S isoscape, generated using the RF regression model that demonstrated the best performance, has been used to calculate predicted values for the environments of the Palaeolithic sites in this research.

To test whether Palaeolithic human populations obtained their prey within the surrounding area of each archaeological site, we generated foraging radii of 1.2 h and 2.5 h around each site, according to the calculation proposed by Marín-Arroyo (2009b), considering the maximum travelling time for ibex and red deer hunting parties using ArcGIS 9.1 Model Builder from ESRI (STable 8). This workflow involves generating a Digital Elevation Model (DEM) suitable for modelling movement through the territory in prehistoric times. Therefore, a new terrain model was built by merging Digital Terrain Models (DTMs) from different sources: LiDAR, EU-DEM, Copernicus, and EMODnet Bathymetric data. Both DTMs were resampled to a similar spatial resolution of 25 meters per pixel, and the data were cropped to elevations below -80 meters relative to the current sea level. The proposed workflow first creates a friction map, also known as a cost surface, from the original DEM (Continental platform + bathymetry). Friction represents how humans ideally move across a rough terrain. For these purposes, the Aragon Mountain Federation algorithm is used in its Python format within the GIS software: Con(“%slope.tif%” <= 0, ((0.6 * 1) * (“%slope.tif%” / 23) +1) / 60, (0.6 * 1) * ((“%slope.tif%” / 11)+1)/ 60). This algorithm produces a map whose pixels indicate the time, in minutes, a human would normally take to traverse that land unit. Once this friction map in raster format (.tif) is available for the research region, a cumulative cost surface is calculated for time zones of 1.2 h and 2.5 h from known sites. This means that each cumulative surface represents the area that can be reached on foot within either 1.2 or 2.5 hours, taking into account the surrounding orography. The third step is to calculate the area in square km and the percentage of each slope type within each time (catchment) area. Therefore, the original DEM is used to calculate slope percentages <30% and >30%. The GIS intersection tools simplify calculating the percentage of each slope class within the Catchment Area. The predicted δ^34^S values within the 2.5 h foraging radius for each site were compared with the observed δ^34^S values for each cultural period and site using boxplots.

### Ecological diversity indices

Two indices were estimated to evaluate the biodiversity of the macro- and micro-mammal communities at the archaeological levels under study. Simpson’s 1/D index combines species richness and evenness within an ecosystem, while Shannon’s H index measures species heterogeneity and their relative richness (Hill, 1973). Thus, both indices quantify the taxonomic diversity by considering richness and evenness. The greater the diversity of ungulate species, the higher the value. To do so, a compilation of published archaeozoological data was assembled (STable 9 and STable 10), including the list of represented species and their MNI in each site and level. Later, we used the “vegan” R package to calculate their values (Oksanen et al., 2013). While micromammals can offer insights into environmental and climatic fluctuations over time (Andrews, 1990), macromammals provide information on the dietary diversity achieved by human groups (Binford, 1981).

## Results and discussion

### Collagen preservation and sample integrity

The stable isotope values for the specimens analysed are reported in STable 3. From the 905 individuals under study, 52 specimens fell outside of the accepted C:N, C:S and N:S ratios and were excluded from analysis. Additionally, 356 samples had %C, %N, and/or %S slightly outside the established margins, but their atomic C:N, C:S, and N:S ratios were within the expected range for well-preserved bone. To test the reliability of such results, we conducted two sensitivity tests at the site and species level. The δ^13^C, δ^15^N, and δ^34^S values of these specimens are not significantly correlated with their %C, %N, or %S, or with their C:N, C:S, and N:S ratios. Similarly, there are no statistically significant differences between the isotopic values of these samples and those obtained with the accepted %C, %N, and %S standards. For both reasons, these samples were included in the analysis. An extended version of the isotopic results, at the site- and species-level, can be found in SText 3.

### ZooMS

ZooMS identification was successful for all samples, and the results are shown in STable 4. The ZooMS species representation is consistent with the archaeozoological associations of these assemblages. All samples were identified as specific ZooMS taxa (*Equidae*, *Capra* sp., *Cervidae/Saiga*, and *Bos/Bison*). Besides, ZooMS has enabled the identification of *Rangifer tarandus*, a taxon that is infrequently represented in the regional archaeological record, especially to the west of the Basque Country. Twenty-four of the 31 samples analysed yielded a different taxonomic identification from that obtained in the zooarchaeological study due to their high degree of fragmentation. 12 of 17 samples previously identified as red deer were found to be another species after proteomic analysis. In particular, eight were *Bos/Bison*, three were *Rangifer tarandus,* and one was *Capra* sp. The other 12 specimens analysed belong to *Bos/Bison* sp. and *Equus ferus*. Six of seven *Bos/Bison* samples were identified as *Equidae*, while six of the seven *Equus ferus* were identified as *Bos/Bison* (*n*= 5) and *Cervidae* (*n*= 1). The *Cervidae/Saiga* category includes several species: *Cervus elaphus, Megaloceros giganteus, Dama dama* and *Saiga tatarica*. The observation of COL111 910 – 934 (2216 m/z) peptide marker excluded *Alces alces*. However, based on the currently available regional faunal information for the Late Pleistocene, an attribution to *Cervus elaphus* is more likely. Therefore, the isotopic values of these samples have been assigned to red deer. *Equidae* also includes several species from the genus *Equus*, although they most likely represent *Equus ferus* (Torres-Iglesias et al., 2024). These results caution against conducting isotopic studies on highly fragmented Palaeolithic faunal assemblages from old excavations, highlighting the need to combine archaeozoology and ZooMS during the sampling phase of future stable isotope studies.

### Isotope results by species

Carbon and nitrogen isotope ratios across all taxonomic groups are consistent with those of ungulates living in C_3_ ecosystems, as documented in Late Pleistocene Europe (Stevens and Hedges, 2004; Stevens et al., 2009; Drucker et al., 2011; Jones et al., 2018; Wißing et al., 2019; Britton et al., 2023a). Differences in feeding behaviour among herbivore species are generally observed across cultures and sites (SFig. 1-8; SText 3). Aurochs/bison, red deer and ibex tend to exhibit similar δ^13^C values, suggesting that these species occupied comparable open to open forest environments (Fig. 2A). Notably, the high variability in red deer δ^13^C values reflects the ecological plasticity of this taxon and its ability to respond to different environmental conditions. In contrast, horses usually display typically lower δ^13^C values, likely reflecting the consumption of poor-quality, low-protein, high-fibre browse (Sponheimer et al., 2003; Jones et al., 2018; Fernández-García et al., 2024). Conversely, reindeer show consistently elevated δ^13^C values, reflecting their typical reliance on lichens (Ben-David et al., 2001) and/or a non-local isozone, given that they are migratory species (Britton, 2018). Regarding δ^15^N values, aurochs/bison exhibit higher values than the other herbivore species (Fig. 2A), consistent with mixed feeders. Mixed-feeder ungulates use both grazing and browsing strategies, incorporating higher-nutrient resources such as short grasses, sedges, forbs, leaves, and shrubs (Merceron et al., 2014). While δ^13^C values show the expected differences among herbivorous taxa, a wide range of δ^15^N values was observed (Fig. 2A), suggesting substantial intra-individual variability that reflects animals procured from distinct microenvironments, as previously reported (Rofes et al., 2015; Jones et al., 2018; 2019).

**Fig. 2.**
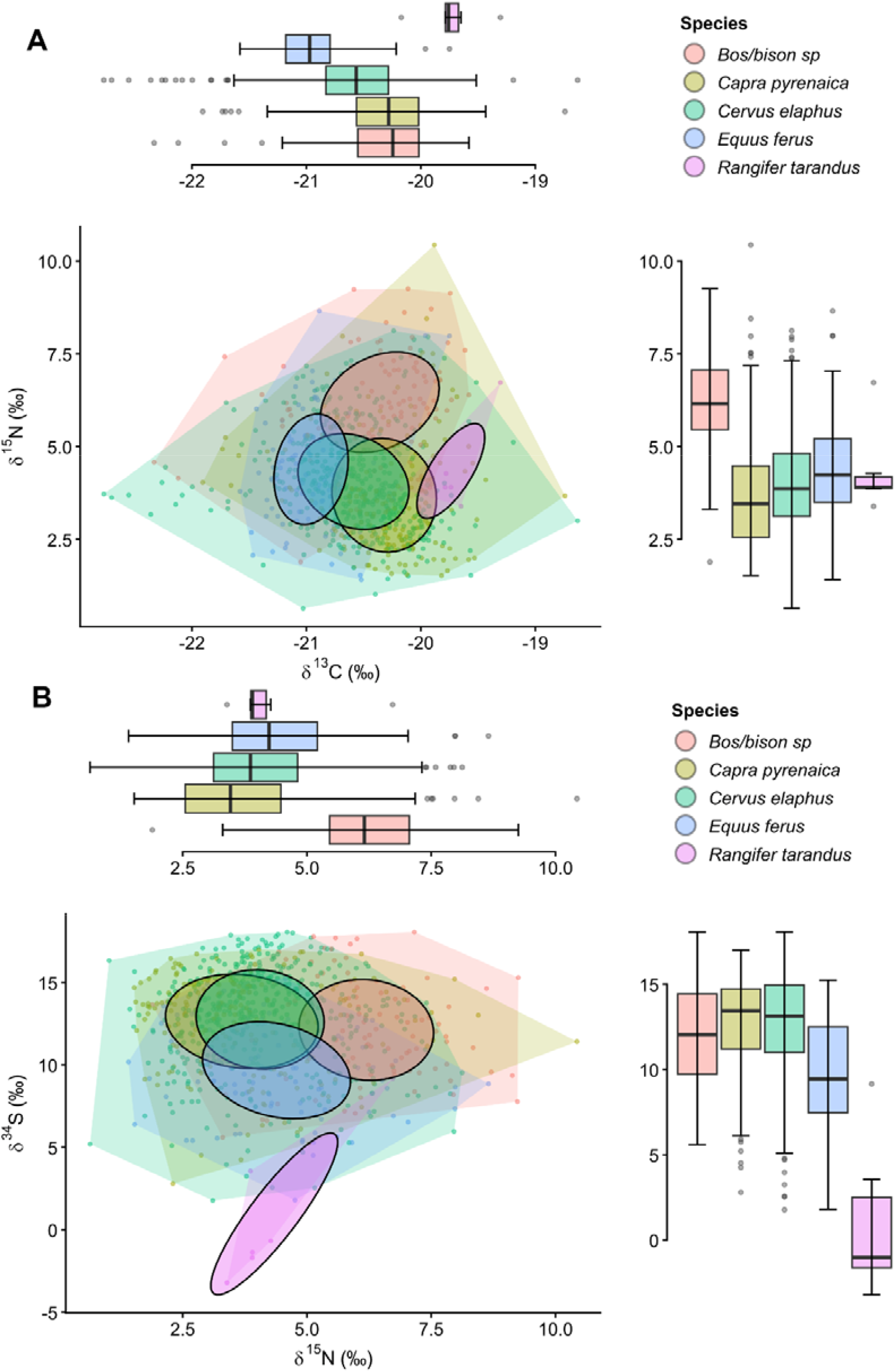
Feeding/spatial niche partitioning between δ13C-δ15N values (A) and δ15N-δ34S va lues (B) for Bos/Bison sp., Capra pyrenaica, Cervus elaphus, Equus ferus and Rangifer tarandus, considering all cultural phases together. Marginal boxplots show the distribution of δ13C, δ15N and δ34S values by species.

Sulfur isotope data show wide ranges in δ^34^S values across all sites, consistent with the trends observed for nitrogen (Fig. 2B). The δ^34^S values of red deer, large bovids, and ibex are not significantly different, suggesting that these taxa coexisted within the same sulfur isozones during most periods (STable 11). Indeed, red deer δ^34^S values only differ from those of aurochs/bison during the Mousterian (*p*= 0.020), indicating that these species usually shared similar δ^34^S niches. Likewise, red deer and ibex do not show statistically significant differences in most sequences, except for the Azilian (*p*= 0.007), suggesting they mostly occupied similar sulfur isozones (isozones= microenvironments with distinct natural or anthropogenic isotopic baselines) (Stevens et al., 2013). In contrast, horses and reindeer indicate occupation of distinct sulfur isozones across cultures (Fig. 2B). Coexistence between horses and other species is rare, with particularly significant differences during the Solutrean and Magdalenian (STable 11), suggesting that horses may have inhabited a location with distinctly lower δ^34^S values. This trend may also have been present during the earliest Upper Palaeolithic periods, although smaller sample sizes likely prevented the detection of statistically significant differences. Although modern wild horses likely have relatively large home ranges and may have undertaken seasonal movements (Schoenecker et al., 2023), the current isotopic evidence does not support long-distance migrations for this species during the Palaeolithic (Barakat et al., 2026). Instead, their physiology and preference for open grassland and steppe environments, which, in mountainous regions, may have been more extensively developed on plateaus, uplands, or exposed slopes, can better explain these lower sulfur values (Sommer et al., 2011). A similar pattern is evident for reindeer, a species whose δ^34^S values are even more negative and show significant differences, suggesting that they likely originated from a different low δ^34^S values environment and that they are the result of a migratory behaviour (Britton et al., 2023b; 2023a; Barakat et al., 2026) (STable 11).

### Niche partitioning of species

Across all time periods, feeding niche partitioning results (δ^13^C-δ^15^N isotopic spaces) show moderate (30–60%) to high (>60%) proportions of total area (TA) overlap among species, except for reindeer (Fig. 2A, STable 12). In contrast, core area (SEAc) results indicate generally low overlap (<30%), although red deer show moderate overlap with ibex and horses, whereas aurochs/bison and reindeer exhibit near-complete partitioning of their core areas (Fig. 2A; STable 12). Spatial niche partitioning results (δ^15^N-δ^34^S isotopic spaces) show a similar pattern at the level of total area, with most species (again excluding reindeer) displaying moderate to high overlap. However, in the core areas, only the ibex-red deer comparison shows high overlap, while ibex–horse and red deer–horse comparisons show moderate overlap, and aurochs/bison and reindeer exhibit low to null overlap (Fig. 2B; STable 12). When examined by cultural phase, both feeding and spatial niche partitioning follow patterns comparable to those observed in the pooled dataset (SFig. 1-8). Overall, total isotopic niche areas consistently show greater interspecies overlap than core areas, suggesting that while species maintain distinct primary resource use and a degree of niche segregation, they occasionally exploit shared resources, reflecting ecological plasticity.

In the Mousterian, aurochs/bison and horses show clear feeding and spatial niche partitioning, with low or null overlap in both total and core areas, contrary to the δ^13^C_diet_ and δ^18^O_mw_ isotope of Axlor and El Castillo (Pederzani et al., 2023; Fernández-García et al., 2024). Similarly, the aurochs/bison-red deer comparison shows no overlap in the core area and moderate overlap in the total area, confirming previous evidence that suggested slightly different isotopic niches between red deer and aurochs/bison (Pederzani et al., 2023). The horse-red deer comparison ranges from low to moderate in the core area and reaches moderate to high overlap in the total area (SFig. 1; STable 12). In the Châtelperronian, feeding and spatial niche partitioning trends are similar to those observed in the Mousterian, with generally low overlap in both isotopic spaces, except for the red deer-horse comparison, which shows moderate to high overlap in both total and core areas of their spatial niches (SFig. 2; STable 12). In the Aurignacian, niche partitioning is reduced compared with Neanderthal-associated cultures, consistent with previous carbonate-isotope evidence, at least in the aurochs/bison and horse comparison (Fernández-García et al., 2024). Both feeding and spatial niches show moderate to high overlap in total areas and low to moderate overlap in core areas across most species comparisons (SFig. 3; STable 12). The only exception is the red deer-horse comparison, which exhibits consistently high overlap in both feeding and spatial niches. In the Gravettian, feeding niche overlap is generally low to moderate in both total and core areas. Aurochs/bison show strong niche partitioning, whereas red deer display moderate to high overlap with ibex and horses (SFig. 4; STable 12). In contrast, the spatial niches of red deer, ibex and horses are highly overlapped, while aurochs/bison remain largely segregated. Previous δ^13^C and δ^15^N isotopic studies had stated a niche separation between horses and red deer after the end of Middle Palaeolithic and initial Upper Palaeolithic cultures (Jones et al., 2018; 2019), but our isotopic data soften this claim and align with δ^13^C_diet_ and δ^18^O_mw_ values of same sites, which exhibit shared ecological niches (Fernández-García et al., 2024).

In the Solutrean, feeding niche overlap among red deer, ibex, and horses is low in both total and core areas, whereas spatial niche overlap is moderate to high, particularly for ibex–red deer and ibex–horse comparisons (SFig. 5; STable 12). The feeding niche partitioning of ibex and red deer in the Solutrean levels of La Riera and Las Caldas was originally proposed by Jones et al. (2020; 2021). In the Magdalenian, ibex and red deer show high overlap in both feeding and spatial niches across total and core areas, also similar to previous δ^13^C and δ^15^N values from the Magdalenian levels of La Riera as a result of the climatic improvement at the end of LGM (Jones et al., 2020). In contrast, niche segregation was maintained in the Magdalenian layers of Las Caldas (Jones et al., 2021) and El Mirón (Geiling, 2020). Horses show moderate overlap in total areas, but null to low overlap in core areas relative to red deer and ibex, similar to the isotopic evidence extracted from teeth of El Otero (Fernández-García et al., 2024). Reindeer exhibit near-complete niche partitioning relative to all other species (SFig. 6; STable 12). In the Azilian, ibex and red deer show high overlap in both total and core areas of their feeding niches, but only moderate overlap in their spatial niches (SFig. 7, STable 12). By contrast, in the Mesolithic, these two species become more clearly partitioned, with low overlap in both total and core areas (SFig. 8, STable 12). Both trends were primarily registered in the Azilian and Meoslithic sequences of La Riera (Jones et al., 2020).

### Climate influence on δ^34^S values

We evaluated the accuracy of MAT and MAP climate estimates derived from the rodent assemblages using the Bioclimatic Model (Royer et al., 2020) against those already accepted from the HadCM3 climate model (Krapp et al., 2021). While MAT and MAP values from the Krapp model closely track the climatic evolution recorded in the NGRIP record (North Greenland Ice Core Project Members, 2004) (SFig. 9), the MAT and MAP estimates from the Royer model show a poorer fit and introduce greater variability (SFig. 10). A direct comparison between models reveals a positive correlation for MAT estimates (*p*= 0.004), whereas MAP estimates predicted by rodent assemblages show no significant correlation with those derived from the general circulation model (*p*= 0.423) (SFig. 11). This suggests that the bioclimatic model may provide a reasonable proxy for temperature but is less reliable for precipitation (Fernández-García et al., 2023). Consequently, MAT and MAP values derived from the Royer model provide substantial uncertainty and should be interpreted with caution.

The limitations of the Bioclimatic Model likely reflect the extremely high-relief, mosaic nature of the landscape. The presence of diverse microhabitats within river valleys has previously been suggested based on stable-isotope and micromammal evidence (Jones et al., 2018; 2019; Fernández-García et al., 2023). Moreover, the complex orography of the Cantabrian Region promotes rapid ecological variation over short distances. As micromammal assemblages capture local environmental signals—unlike general circulation models—they are also more susceptible to taphonomic biases, excavation issues, and taxonomic uncertainties that may distort species representation (Álvarez-Vena et al., 2021). In addition, 39 of the 111 of the archaeological levels analysed in this study lack micromammal data. For these reasons, we rely on paleoclimate reconstructions from the HadCM3 climate model (Krapp et al., 2021) to assess the influence of MAT and MAP estimates on δ^34^S values in bone collagen.

Statistical analyses did not reveal any correlation between δ^34^S values and MAT estimates in any species (red deer, horses or ibex), with the exception of aurochs/bison (*p*= 0.029) (STable 13). In contrast, MAP estimates show positive correlations with δ^34^S values in aurochs/bison (*p*= 0.031) and horses (*p*= 0.026), possibly reflecting their aforementioned dietary ecology, physiology and the specific isotopic niches they occupy (Sponheimer et al., 2003; Merceron et al., 2014). Red deer and ibex—the most abundant taxa in this study—do not show any relationship between δ^34^S values and climatic variables, suggesting that factors such as temperature, aridity, vegetation cover, and atmospheric CO₂ are not primary drivers of sulfur isotopic variation during the Palaeolithic (Jones et al., 2018), unlike what has been proposed for carbon (Stevens and Hedges, 2004; Drucker et al., 2008; Stevens et al., 2025) and nitrogen (Amundson et al., 2003; Richards and Hedges, 2003; Stevens and Hedges, 2004; Stevens et al., 2008; 2014; Drucker et al., 2011). These results indicate a limited, indirect influence of climate on the sulfur isotope system and highlight interspecies variability in responses to climatic and environmental fluctuations in this region. To further explore this relationship, MAT and MAP estimates were correlated with mean δ^34^S values from archaeological levels grouped by MIS, with red deer as the most representative species (analyses restricted to MIS 3–1 due to limited sample sizes for MIS 5 and 4). While no correlation between δ^34^S values and MAT estimates was observed for MIS 1, significant correlations were identified for MIS 2 (negative; *p*= 0.002) and MIS 3 (positive; *p*= 0.003) (STable 14), indicating that the relationship between temperature and sulfur isotopes was not linear through time. For MAP estimates, only MIS 3 shows a significant positive correlation with δ^34^S values (*p* < 0.001), suggesting a weak relationship between δ^34^S values and precipitation (STable 14).

Site-level correlations between MAT, MAP, and δ^34^S remain inconsistent. Some negative correlations between MAT and δ^34^S are observed at El Mirón (*p* < 0.001) and Las Caldas (*p*= 0.009); while positive correlations between MAP and δ^34^S are identified at Aitzbitarte III (*p*= 0.044), El Castillo (*p*= 0.036) and Las Caldas (*p* < 0.001) (STable 15). These patterns suggest that local sulfur values are influenced by site-specific environmental and climatic conditions operating at a microregional scale. Overall, the influence of climatic factors on the sulfur isotope system shows distinct dynamics across species and climatic periods, suggesting that they exert variable, indirect and generally limited control over sulfur isotope variability and demonstrating that palaeoclimatic changes are not a primary driver of δ^34^S variations.

### Correlations with δ^13^C and δ^15^N isotope values

To assess changes in isotope values throughout the Palaeolithic in the Cantabrian Region and to test potential links between sulfur isozones and the carbon and nitrogen values isotopic niches, we performed statistical correlations between δ^13^C/δ^15^N, and δ^34^S values of faunal species from the archaeological levels studied. No significant correlations between δ^13^C and δ^34^S values are observed for aurochs/bison, horses and ibex, whereas red deer shows a significant negative correlation (*p* < 0.001) (STable 16). In contrast, all species exhibit a statistically significant negative correlation between δ^15^N and δ^34^S values, indicating a strong link between nitrogen and sulfur isotope values within this dataset, as previously suggested by the large inter-individual variability observed in both isotopes. This pattern is consistent across the chronological sequences of each taxon (Fig. 3, SFig. 12-14), with higher δ^15^N values generally associated with lower δ^34^S values.

**Fig. 3.**
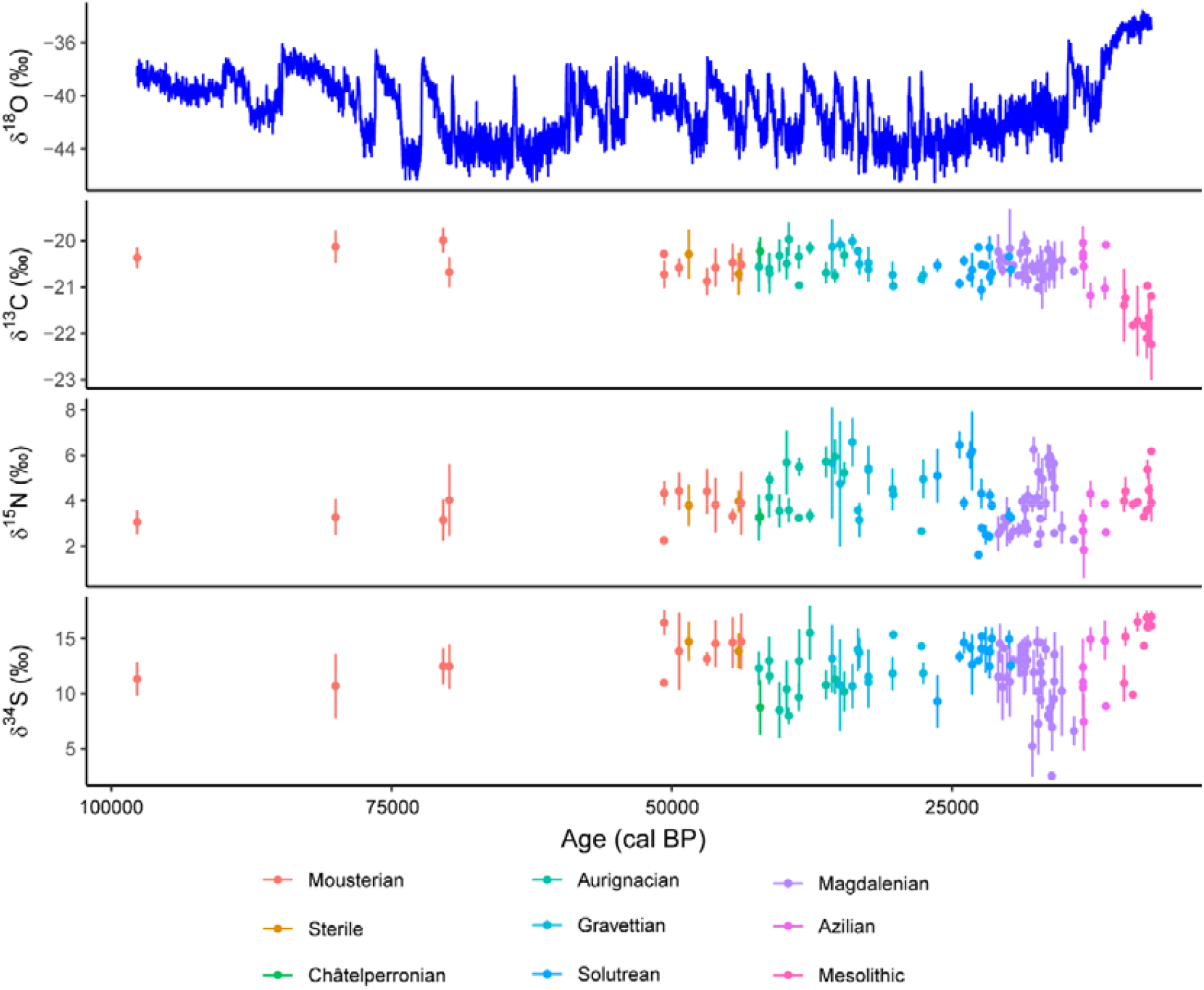
Mean δ13C, δ15N and δ34S values (error bars show 1σ) of red deer for each archaeological level against NGRIP δ18O isotopes (Rasmussen et al., 2014).

Two particularly marked intervals show pronounced decreases in δ^34^S values among herbivores, coinciding with increases in δ^13^C and δ^15^N values: the first between ca. 42–38 ka BP and the second between ca. 17–15 ka BP (Fig. 3). Following these episodes, δ^34^S values increase and subsequently stabilise, whereas δ^13^C and δ^15^N values decrease. The decreases in δ^34^S values are coeval with Heinrich Stadial 4 (HS4) and Heinrich Stadial 1 (HS1), two episodes characterised by extremely cold and arid conditions linked to a weakening of the Atlantic Meridional Overturning Circulation (AMOC) (Hemming, 2004). At a regional scale, both events also coincide with relevant deglaciations in the Cantabrian Mountains, marked by glacier retreat, particularly during HS1, which represents the most extensive deglaciation phase in the region (Moreno et al., 2010; Frochoso et al., 2013; Rodríguez-Rodríguez et al., 2017; Serrano et al., 2017) and with an increase in net primary productivity (NPP) for the first episode of δ^34^S values decrease (Vidal-Cordasco et al., 2022).

Two hypotheses related to the indirect impact of environmental changes are proposed to explain the lower sulfur values observed in both episodes. First, glacier melting—combined with reduced precipitation and rising temperatures—may have altered the water cycle (Stevens et al., 2023). In this scenario, increased weathering and oxidation of ^34^S depleted bedrock sulfide minerals, and release of ^34^S depleted glacial waters stored in ice masses over millennia would result in ^34^S depleted streamwater sulfate being transported to isozones downstream (James et al., 2026). Thus, animal herds would have remained within similar territories, and hunter-gatherer groups would have continued to exploit these areas under changing environmental conditions, particularly with respect to water availability, without substantial shifts in mobility patterns. Alternatively, the retreat of glaciers may have exposed new landscapes and grazing areas, allowing ungulates to expand their ranges further inland, up the valleys, and away from coastal zones (Straus, 2015). In this case, human groups may have followed these herds over greater distances during hunting parties, leading to adjustments in hunting strategies and mobility behaviours in response to shifting environmental conditions.

### Cultural trends

Sulfur isotope values show considerable variability across all species (Fig. 4, STable 3, SText 3). This pattern is evident within archaeological levels and across cultural phases of each site, suggesting that these herbivores were hunted across distinct isozones. Mousterian sites display generally high δ^34^S values, although the inland sites of Llonín and Axlor, yield slightly lower values than El Castillo, Covalejos, and Amalda (Fig. 4A). Meanwhile, the only Châtelperronian level analysed, from Labeko Koba—the site furthest from the coast—shows notably low sulfur values (Fig. 4B). Aurignacian sites (La Viña, El Castillo, Covalejos, and Labeko Koba) exhibit higher δ^34^S values, but slightly lower than those observed in the Mousterian. An exception is Aitzbitarte III, which shows markedly higher δ^34^S values (Fig. 4C). Gravettian sites (La Viña, Llonín, Bolinkoba, and Amalda) follow a similar pattern, although Aitzbitarte III displays lower values and greater standard deviation than during the Aurignacian (Fig. 4D). Solutrean sites again show relatively high δ^34^S values, although La Viña yields slightly lower values than Las Caldas, La Riera, El Mirón, and Amalda. Interestingly, this phase is characterised by reduced inter-individual variability (Fig. 4E). Magdalenian sites, both coastal and inland, display substantial and high standard deviations in sulfur values. Higher δ^34^S values are observed at La Riera, Altamira, El Juyo, and El Mirón, whereas Las Caldas and La Viña, located more inland, show lower values, below those recorded for the Solutrean (Fig. 4F). Azilian sites exhibit two contrasting patterns: the coastal site of La Riera shows high δ^34^S values, while the inland site of El Mirón shows lower δ^34^S values, both with considerable variability (Fig. 4G). Finally, Mesolithic sites, including La Riera, El Mazo and El Toral III, located along the present-day coastline, yield the highest δ^34^S values of the entire cultural sequence, exceeding those of El Mirón, located in the middle Asón Valley. These sites also show the lowest sulfur inter-individual variability (Fig. 4H).

**Fig. 4.**
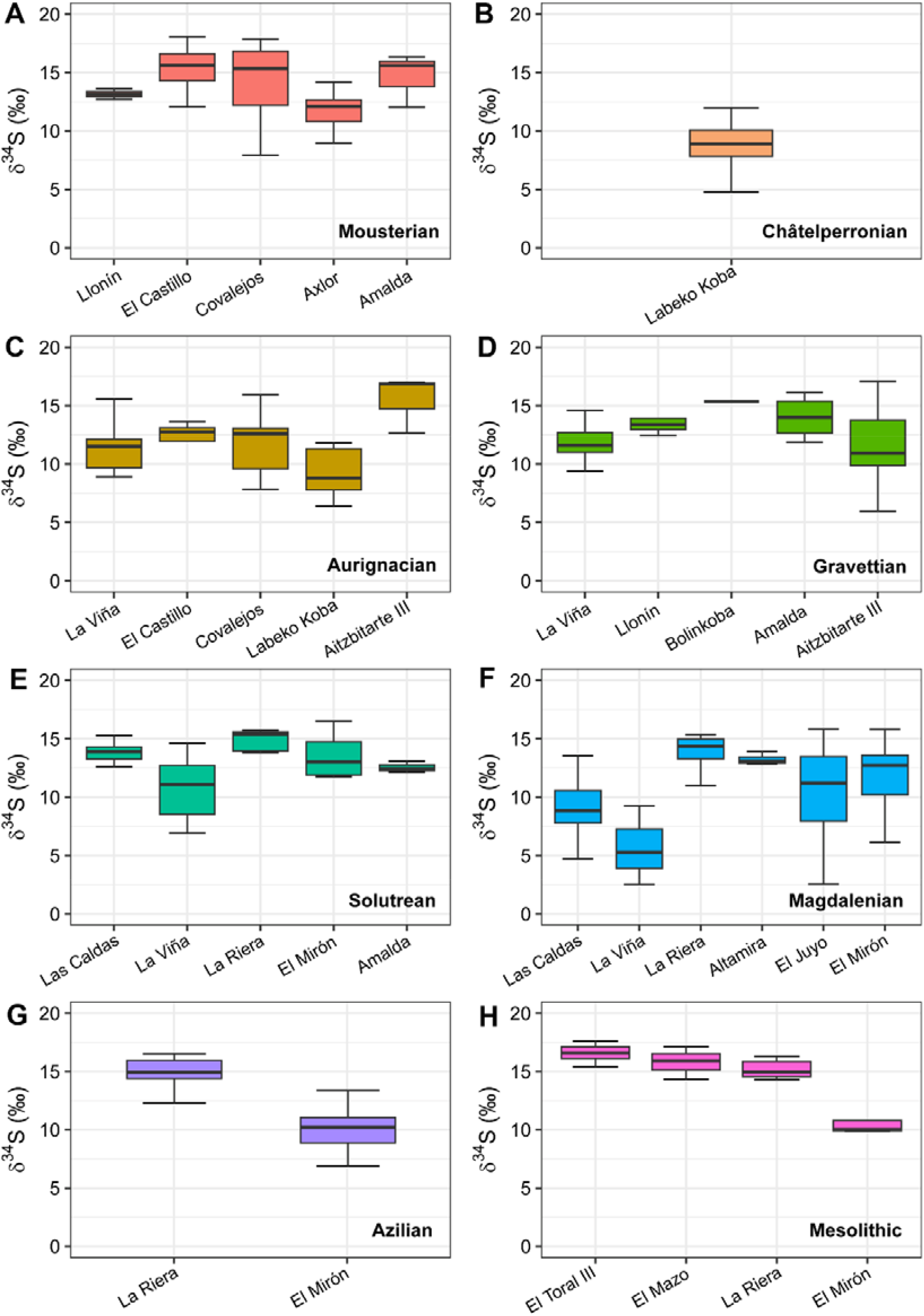
Boxplots of the δ³⁴S values grouped by cultural phases and considering only Cervus elaphus individuals.

Given the aforementioned variability, the inclusion of individual δ^34^S values from different sites limits the reliability of diachronic comparisons for determining how sulfur values changed across Palaeolithic cultures. To address this, we analysed mean sulfur values (δ^34^S*m*) and sulfur standard deviation (δ^34^S*sd*) for each archaeological level and cultural phase, recognising that higher/lower δ^34^Sm values indicate areas more enriched/depleted in ^34^S, and that higher/lower δ^34^S*sd* values imply greater/lesser inter-individual variability. Our results indicate that red deer from the Mousterian, Solutrean, and Mesolithic periods were more frequently hunted in isozones with higher δ^34^S values, likely near coastal environments. In contrast, red deer from Aurignacian, Gravettian, Magdalenian and Azilian were exploited in more ^34^S depleted areas (Fig. 5A), with statistically significant differences in several cases (STable 17), suggesting that these human groups hunted prey in both coastal and inland areas and were thus more mobile. Red deer also show relatively high variability, with higher δ^34^S*sd* values across all cultures except the Solutrean and Mesolithic, when significantly lower δ^34^S*sd* indicates more restricted human mobility (Fig. 5B, STable 17). Instead, aurochs/bison display relatively consistent δ^34^S*m* and δ^34^S*sd* results across cultural phases, with no statistically significant differences (SFig. 15, STable 17), although the small sample size warrants caution. Horses show lower δ^34^S*m* values than other species, particularly during Aurignacian and Magdalenian (SFig. 16A), with statistically significant differences in the latter (STable 17). Notably, Gravettian horses display the greatest dispersion, with higher δ^34^S*sd* values (SFig. 16B), also statistically significant (STable 17). Ibex show slightly lower δ^34^S*m* values during the Magdalenian (SFig. 17A), which differ significantly from those of the Solutrean and Azilian (STable 17). Overall, ibex exhibit the lowest variability, with consistently low δ^34^S*sd* values, and limited differences between cultures, although slightly higher variability is observed during the Solutrean and Magdalenian (SFig. 17B).

**Fig. 5.**
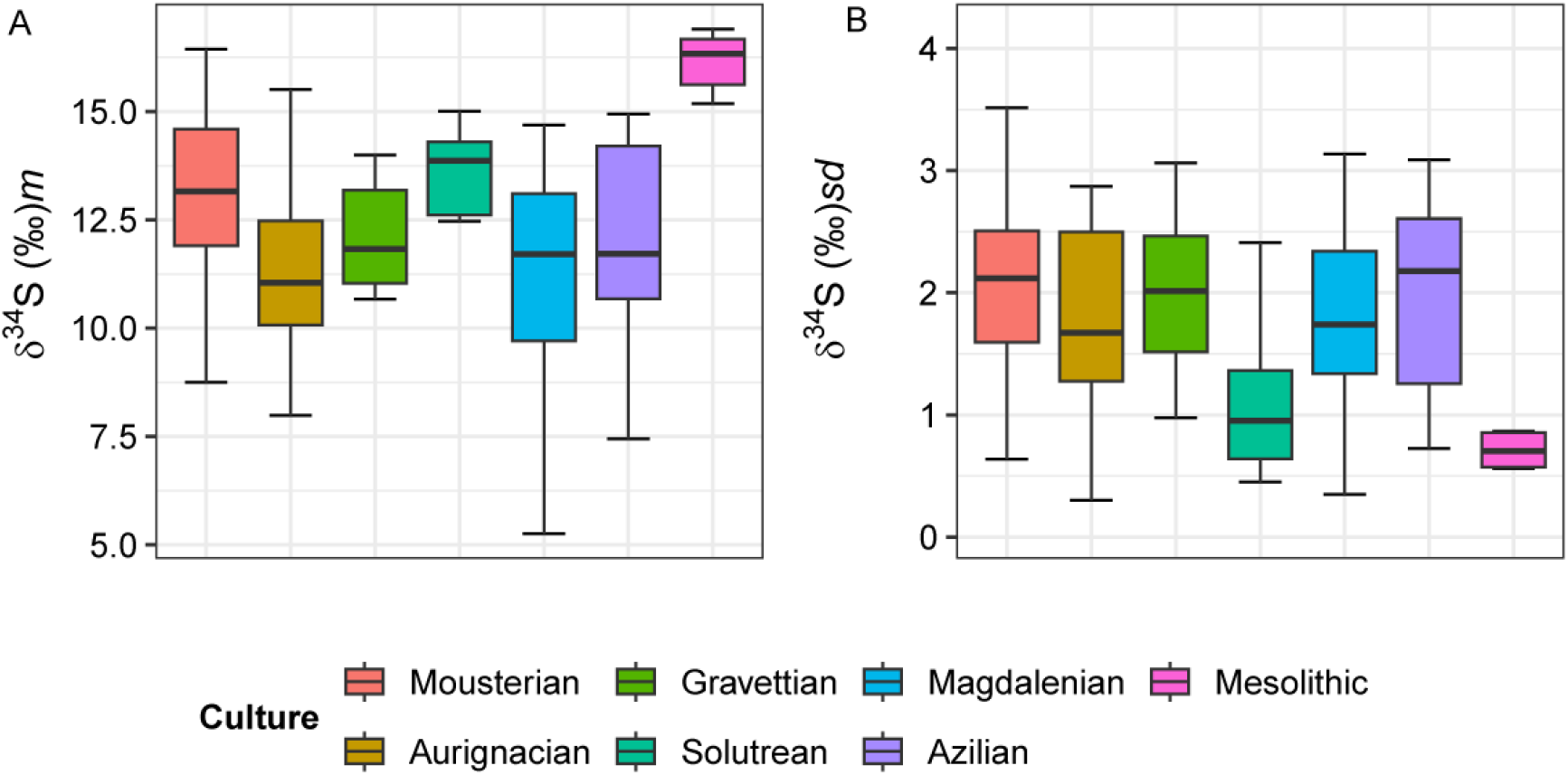
Boxplots showing the mean (A) and the standard deviation (B) of δ³⁴S values for the archaeological levels, grouped by cultural phases and considering only Cervus elaphus individuals.

### Overlaying the sulfur isotopic values and catchment areas of the sites

The integration of site locations and their respective catchment areas into the sulfur isoscape of the Cantabrian Region reveals distinct sulfur isozones within 1.2 and 2.5 hours of each site. Overall, the isoscape displays two consistent spatial trends: 1) coastal areas exhibit elevated δ^34^S values than inland regions; and 2) western coastal areas show elevated δ^34^S values in comparison to eastern ones (Fig. 6). These patterns highlight substantial spatial heterogeneity in sulfur isotope distribution across the region, which likely contributes to the wide variability observed in the sulfur isotope values of sampled ungulates. When overlaying the predicted δ³⁴S values within the 2.5-hour catchment areas to those measured in animal bone collagen of animals hunted at each site, the Holocene isoscape model developed by González-Rabanal et al. (2025) performs reasonably well (STable 18). Most measured values fall within the predicted ranges, and both mean and maximum values show a positive correlation with model expectations. However, minimum values do not correlate, suggesting that the lowest δ³⁴S values derive from animals hunted beyond the defined catchment areas, likely in more inland environments.

**Fig. 6.**
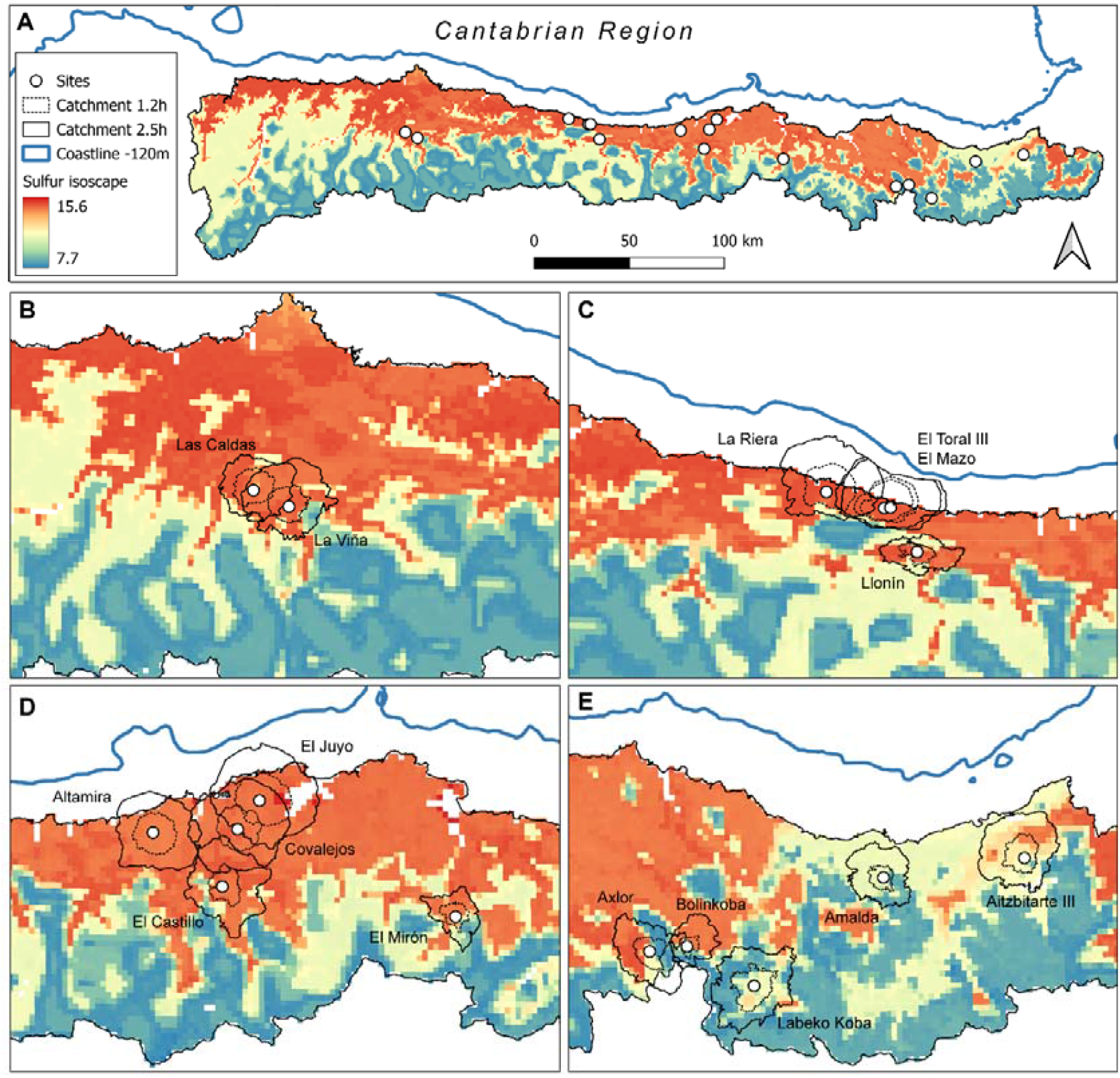
Sulfur isoscape model reconstruction for Northern Iberia (González-Rabanal et al. 2025): A) Spatial distribution of the predictive sulfur isotope composition (δ34S) across the Cantabrian Region. B) Spatial distribution of the sulfur isotope composition (δ34S) in the vicinity of the archaeological sites analysed in this research (white dots) surrounded by the catchment areas of 1.2 h and 2.5 h around each site (black lines).

A comparison across cultural phases and sites indicates that archaeological samples consistently exhibit a broader range of δ³⁴S values (*sd*= 2.5‰) than those expected from local catchment areas (*sd*= 1.2‰), most of them even over the ∼2‰ intra-individual variability commonly observed in modern studies of herbivores from a single population (Nehlich, 2015). This discrepancy indicates that the isotopic composition of hunted animals does not fully explain the local environmental signatures alone (SFig. 18, SFig. 19), suggesting that the current isoscape serves as a framework but does not capture all the real variability and bioavailability of sulfur isotopic systems within the catchment areas, or alternatively, that human mobility extended beyond the estimated foraging territories.

For the Mousterian, measured and predicted δ³⁴S values align closely, implying limited mobility and a stronger reliance on local resources; statistical comparisons do not show significance (STable 19). In contrast, the Châtelperronian shows noticeable dispersion, with several values lower than expected and this was statistically significant (STable 19). This pattern suggests that the subsistence practices of the Châtelperronian groups were not strictly confined to the immediate catchment area and may have involved exploiting environments far inland. This evidence becomes more pronounced in the Aurignacian, when measured values frequently diverge from predictions, suggesting that early Anatomically Modern Humans (AMHs) engaged in longer-distance movements, particularly toward inland low δ^34^S isozones and probably following the herds in a N-S direction. This trend is consistent across most Aurignacian sites except Aitzbitarte III (SFig. 18, SFig. 19).

For the Gravettian and Solutrean, measured and predicted values generally converge again, pointing to reduced mobility and a focus on nearby hunting territories, although La Viña remains an exception with lower measured δ³⁴S values. The general pattern of a broader range of measured δ³⁴S values than predicted for the catchment areas is particularly pronounced in Magdalenian contexts, with all sites showing significant differences (STable 19). This increased sulfur variability may reflect greater mobility, expanded territorial ranges, especially into low δ³⁴S isozones, or more diverse subsistence strategies. The presence of outliers further supports the existence of individuals with distinct isotopic signatures, potentially reflecting long-distance movement to inland territories for faunal procurement without discarding inter-regional contacts (SFig. 18, SFig. 19). Finally, in the Azilian and Mesolithic periods, measured sulfur isotope values are closer to those predicted by the catchment areas. In fact, many of the sites do not show statistically significant differences (STable 19). However, they also exceed, in some cases, the local environment, particularly towards higher δ³⁴S isozones, which are probably related to territories even closer to the early Holocene coastline (Fig. 6; SFig. 18, SFig. 19).

### Ecological diversity trends

Macromammal and micromammal assemblages show a significant negative correlation in Simpson’s 1/D (*p*= 0.003) and Shannon’s H (*p*= 0.002) indices (SFig. 20), indicating that the trends in ecological diversity of both communities followed divergent trajectories in the Cantabrian Region. This correlation could be explained by seasonal behaviour, in which predators and humans could not have occupied the caves simultaneously (Pokines, 2000). So, the longer human occupation of these sites, the higher the expected macrofaunal diversity, whereas avian visits become less frequent, and therefore lower microfaunal diversity. However, this evidence would be expected if both communities were governed solely by environmental and ecological factors. However, their differing site formation processes help explain this discrepancy: micromammals, typically accumulated by avian predators, reflect local environmental conditions (Álvarez-Vena et al., 2021), whereas macromammals primarily represent human subsistence choices (Marín-Arroyo, 2013). Alternatively, the observed correlation may suggest that during periods of reduced micromammal diversity—likely linked to harsher climatic conditions—hunter-gatherers adjusted their mobility and subsistence strategies, thereby increasing their hunting diversity according to species (Cuenca-Bescós et al., 2012).

The chronological changes in the representation of macromammal communities reveal fluctuating diversity throughout the Palaeolithic, reflecting shifts in both richness and evenness (Fig. 7). However, these changes may differ slightly at coastal or montane sites, with changes in biodiversity patterns. Mousterian levels are characterised by relatively lower Simpson and Shannon values, whereas the Aurignacian shows a marked increase, although previous studies suggest more stable average species diversity across these periods (Marín-Arroyo and Sanz-Royo, 2022). Our results indicate that lower species diversity seen in the Late Mousterian was contemporaneous with a significant decrease in NPP, while higher species diversity in the Aurignacian coincided with an increase in NPP (Vidal-Cordasco et al., 2022). Thus, the increase during the Aurignacian may indicate that AMH exploited a variety of ungulate resources, requiring greater mobility and wider foraging ranges than Neanderthals. High diversity persists into the Gravettian but declines during the Solutrean and Lower Magdalenian, a trend commonly associated with increased specialisation in red deer hunting driven by economic profitability (Freeman, 1973; Straus, 1977; Marín-Arroyo, 2009b; Portero et al., 2024; 2026). From the Middle Magdalenian onwards, species diversity increases again and remains relatively stable into the Azilian. This diversification reflects the inclusion of previously less-exploited taxa such as wild boar and roe deer, alongside a reduced dominance of red deer and ibex. These changes are consistent with progressive reforestation and more temperate climatic conditions, which are likely to alter species availability (Marín-Arroyo, 2013). In the Mesolithic, coinciding with intensified use of littoral environments (Gutiérrez-Zugasti et al., 2011), faunal assemblages show low species diversity, dominated by a few taxa, a pattern that matches previous findings (Portero et al., 2022). Notably, Simpson/Shannon indices show a strong negative correlation with sulfur isotope values (*p* < 0.001), with higher diversity values associated with lower sulfur values (SFig. 21). No comparable correlation is observed with carbon or nitrogen isotopes; therefore, the increase in diversity is interpreted as resulting from the expansion of catchment areas and then greater human mobility. Lower prey abundances around some settlements, due to previous intense exploitation or climate change, would force human groups to travel relatively longer distances – particularly to inland territories with lower sulfur isotope values – to procure prey.

**Fig. 7.**
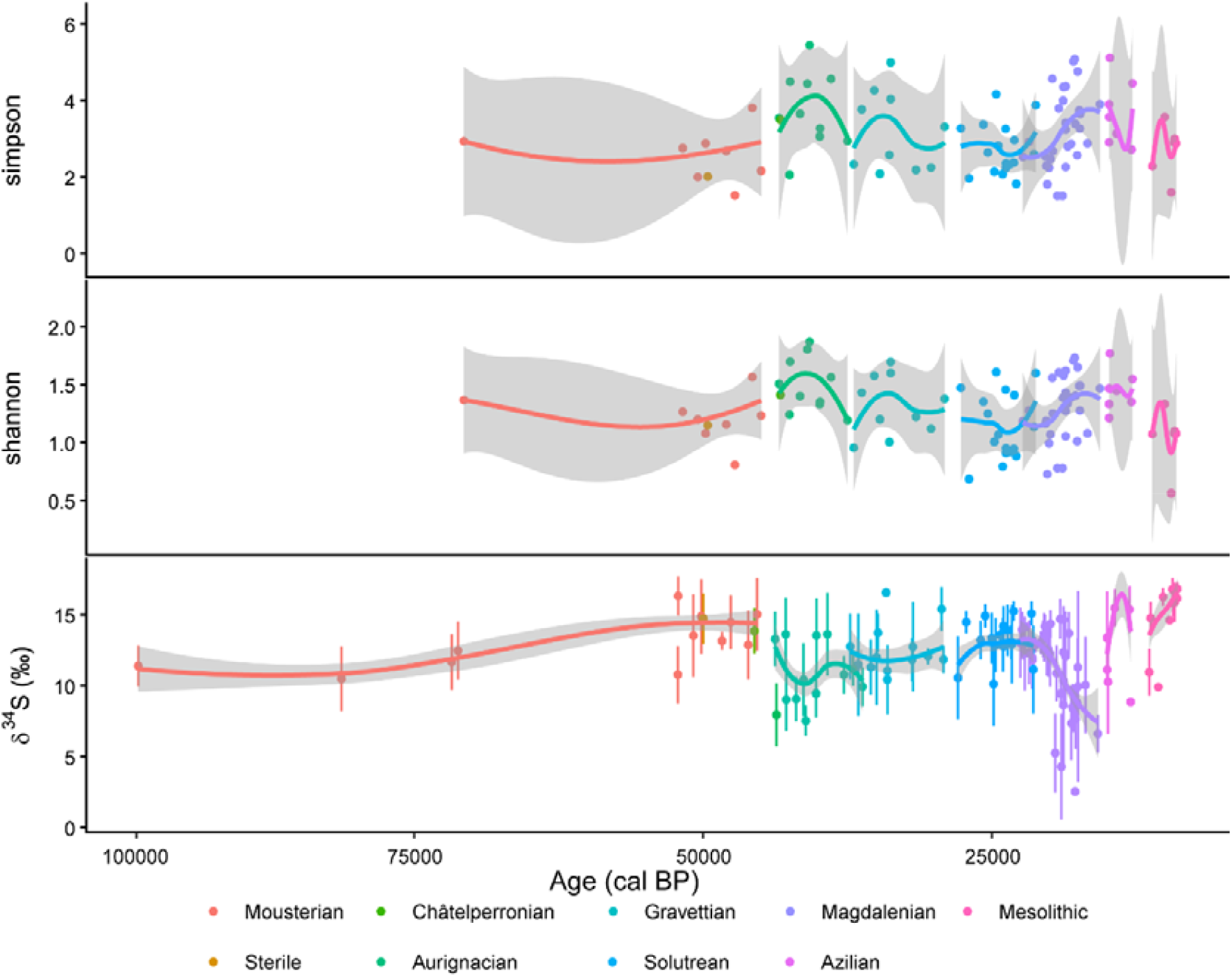
Simpson’s 1/D and Shannon’s H and δ34S values of macromammals for each archaeological level.

In contrast, the trend in micromammal species shows a smoother, but completely inverse trajectory. High diversity is observed during the Mousterian, followed by a decline in the Aurignacian. Diversity increases again in the Gravettian, stabilises during the Solutrean, and decreases through the Magdalenian. By the Azilian and Mesolithic, micromammal diversity converges with macromammal patterns for the first time in the sequence (SFig. 22). Unlike macromammals, micromammal diversity shows a positive correlation with sulfur isotope values (*p*= 0.015), with higher diversity associated with higher δ³⁴S values (SFig. 23). This positive relationship suggests that micromammal communities were more diverse under conditions of stronger marine influence (higher δ³⁴S values), possibly because these environments supported a greater diversity of ecological niches or microhabitats for small mammals (i.e. forest, scrubland, meadows, wetlands, dunes, etc.) (Álvarez-Vena, 2024; González-García et al., 2024) Additionally, we also find significant correlations with carbon and nitrogen isotopes, indicating that climatic and environmental factors were the primary drivers of shifts in micromammal biodiversity.

### Logistical vs residential mobility behaviour

The ungulates analysed are associated with human hunting activities in the vicinity of the archaeological sites and beyond; therefore, their sulfur values result from the local signal from which they were procured and ultimately constitute potential indicators of their mobility behaviours. The range of sulfur isotope values of 48 archaeological levels exceeds the inter-individual variability observed in modern studies (*m*= 1.9‰), while the range of the 29 levels exceeds the inter-individual variability reported in archaeological studies (*m*= 2.4‰) (Nehlich, 2015). Thus, exploitation of a wide variety of isotopically distinct habitats by both animals and humans throughout the Cantabrian Palaeolithic seems likely. All evidence shows that coastal and inland regions were closely linked within a common forager settlement-subsistence system. This territory was exploited through logistical movements, although this model would not limit residential movements between sites for all-season occupation, as proposed by Marín-Arroyo et al (2023). Thus, the whole of the narrow Cantabrian strip between the Palaeolithic shore and the front ranges of the Cordillera would have been used by Upper Palaeolithic foragers to establish residential base camps from which logistical hunting parties would be organised. Despite significant environmental changes induced by abrupt and continuous climatic oscillations between MIS 5 and MIS 1, human groups did not substantially modify their hunting territories, indicating that these territories continued to meet their needs for most of the cultural sequence.

The integration of sulfur isotope records, catchment area modelling, and ecological diversity indices provides a robust framework for assessing mobility strategies among Palaeolithic hunter-gatherers in the Cantabrian Region. Overall, the evidence indicates that mobility was dynamic and flexible over time, combining elements of both residential and logistical systems rather than conforming strictly to a single model. In general terms, the wide dispersion of δ³⁴S values across sites, species, and cultural phases suggests that hunter-gatherers exploited multiple sulfur isozones, including both coastal (high δ³⁴S values) and inland (low δ³⁴S values) environments. This pattern is further reinforced by a partial mismatch between measured isotopic values and those predicted within local catchment areas of some cultures, indicating that prey was often procured beyond the immediate surroundings of the sites. Such evidence points to the recurrent use of logistical mobility, involving targeted forays into distant areas to acquire resources. However, periods characterised by low isotopic variability and strong correspondence with local isoscapes suggest episodes of reduced mobility, more consistent with localised exploitation and potentially more residentially stable settlement systems.

When examined diachronically, mobility strategies show some cultural variability. This research is particularly valuable for discerning subsistence strategies during the Middle-Upper Palaeolithic transition. A recent study emphasises that Neanderthals were strongly influenced in their hunting decisions by the topographic setting and surrounding environment of each site, in contrast to Anatomically Modern Humans (AMH). This would have implied longer travel distances for AMH and greater reliance on logistical mobility (Marín-Arroyo and Sanz-Royo, 2022). Our sulfur isotope results suggest substantial heterogeneity in the hunting areas used by both Neanderthals and AMH in the vicinity of each site, resulting in high inter-individual variability. This is evident in all Neanderthal-associated technocomplexes (Mousterian and Châtelperronian), as well as in the earliest AMH cultures (Aurignacian and Gravettian) (Jones et al., 2018; Pederzani et al., 2023). However, the close correlation between measured and predicted δ³⁴S values, and the lower species diversity during the Mousterian compared to the Châtelperronian, Aurignacian, and Gravettian, could indicate a major reliance on local resource exploitation by Neanderthals, consistent with a strategy of greater residential mobility, in which hunting activities were limited to short-distance logistical movements. These periods show increased variability in sulfur isotope values, frequent deviations from local isoscape predictions and higher species diversity indices. This evidence suggests a shift toward enhanced logistical mobility, with hunting excursions extending into inland territories with low δ³⁴S values. In the Aurignacian, this pattern becomes more pronounced, indicating that early AMH likely operated over larger territories, possibly reflecting changes in social organisation, subsistence efficiency, or population dynamics (Marín-Arroyo and Sanz-Royo, 2022).

During the Solutrean, a return to closer alignment between measured and predicted δ³⁴S values and lower prey diversity suggests a reduction in mobility range and a stronger focus on local environments. This pattern, together with lower isotopic variability in δ³⁴S values (especially focused on coastal areas), is consistent with more structured settlement systems, potentially combining residential stability with shorter-range logistical forays motivated by the harshest climate conditions during the LGM (Straus, 2015; Geiling et al., 2025). Instead, the Magdalenian (especially from the Middle Magdalenian) is characterised by high isotopic variability and significant divergence from local catchment predictions, indicating a renewed expansion of mobility. The exploitation of multiple isozones and the presence of isotopic outliers strongly support intensive logistical mobility, with long-distance hunting trips into inland areas. This is further corroborated by increased faunal diversity from the Middle Magdalenian onwards, suggesting broader territorial use and more flexible subsistence strategies (Straus, 2005). It is precisely at this moment that we identified reindeer individuals with negative sulfur isotope values, suggesting either occasional reindeer incursions into the Cantabrian Region as part of annual migration routes or trade exchanges between hunter-gatherers from very distant places (Gómez-Olivencia et al., 2014; Lefebvre et al., 2021). In the Azilian and Mesolithic, measured δ³⁴S values tend to converge again with local isoscape predictions, and isotopic variability and mammal diversity decrease. This pattern suggests a reduction in mobility scale, with subsistence strategies focused on nearby environments. In the Mesolithic, the strong exploitation of coastal resources and high δ³⁴S values indicate a more restricted and residential mobility system, likely linked to high demographic pressure and territorial behaviour (Arias, 2005), although longitudinal movements along the coast cannot be ruled out.

In sum, the data indicate that Cantabrian hunter-gatherer mobility cannot be described by a single model; rather, it fluctuated between logistical and residential strategies depending on cultural, environmental, and subsistence conditions. Periods of low prey availability, climatic stress, or increased competition appear to have encouraged wider foraging ranges and logistical mobility, whereas resource abundance and environmental stability favoured more localised, residential systems. This variability highlights the adaptability of human groups to climate and demographic conditions, and supports a model of flexible mobility systems, in which logistical and residential components were combined and adjusted over time.

## Acknowledgments

This research began in 2013 and has been made possible by various funding bodies awarded to A.B.M-A. including those from the Marie Skłodowska-Curie Career Integration Grants (CIG 2012- 322112), the Spanish Ministry of Science, Innovation and Universities (refs: PID2021-125818NB-I00; HAR2017-84997-P; HAR2012-33956) and the European Research Council through a Consolidator Grant to the Subsilience project (Ref 818299). All these projects covered analytical expenses and research contracts with B. G-R., J.R.J., M.V-C., L.A.P., L.T.I., M.F-G., H.R., A.S-R., and G.T. Additional MSCA-IF funding (ref: MSCA-IF-2014-656122) was obtained by J.R.J. between 2015 and 2017. During the writing and review of this paper B.G-R was funded by MCIN/AEI/10.13039/501100011033 - the European Union «NextGenerationEU»/PRTR (JDC2022-048798-I) and the Asturian Agency for Science, Business Competitiveness and Innovation of the Principality of Asturias – SEKUENS (IDE/2025/000919); L.T-I was funded by a Marie Skłodowska-Curie grant (HORIZON-MSCA–2023–PF–01; Ref. 101148342); M.V-C was funded by a Marie Skłodowska-Curie grant (HORIZON-MSCA–2024–PF-01; Ref. 101207747); M.F-G was funded by a Marie Skłodowska-Curie grant (HORIZON-MSCA–2025–PF-01; Ref. 101205968); and R.S. was funded by the Leverhulme Trust (RPG-2021-254). We also gratefully acknowledge the laboratory work conducted by G. Terlato during this research. We acknowledge the curators and institutions from the Museo Arqueológico de Asturias (MAA), the Museo de Prehistoria y Arqueología de Cantabria (MUPAC), the Museo de Arqueología de Bizkaia (Arkeologi Museoa) and the Centro de Colecciones Patrimoniales de la Diputación Foral de Gipuzkoa (Gordailua Center) and the Basque Government for access and permissions to sampling archaeological materials deposited at these sites.

## Author contributions

Project conception: A.B.M-A. and B.G-R. Funding acquisition, project administration and supervision: A.B.M-A. Sampling and collagen extraction analysis: A.B.M-A., R.S., H.R., J.J., L.A.P., B.G-R., L.T-I., A.S-R., and M.F-G. Stable isotope analysis: A.B.M-A., B.G-R and J.J. ZooMs analysis: L.A.P. and L.T-I. Bayesian age modelling analysis: B.G-R. and A.B.M-A. Palaeoclimatic reconstruction analysis: M.V-C. and A.A-V. Isoscape mapping analysis: M.V-C. and J.G-S. Ecological diversity analysis: B.G-R. and M.V-C. Visualisation: B.G-R. and M.V-C. Resources and facilities: A.B.M-A., M.P.R., R.S., and T.O. Excavation, access and provision of samples: J.A., K.M., M.S.C-R., D.C-S., M.R.G.M., I.G-Z., P.F., M.D.L.R., and L.G.S. Writing – original draft: B.G-R., A.B.M-A., and J.J. Writing – review & editing: all authors revised and approved the manuscript.

## Data availability

All data needed to support the conclusions of the paper are presented in the main manuscript and the Supplementary Information: SText 1-3, SFigure 1-23, STable 1-19, and SCode 1-6.

